# Not so cold after all: tumor infiltrating CD8+ T cells in EBV-positive Burkitt lymphoma are quiescent, not exhausted

**DOI:** 10.64898/2026.04.15.718702

**Authors:** Catherine S. Forconi, Cliff I. Oduor, Priya L. Saikumar, Zachary J. Racenet, Gavin Fujimori, Viriato M’Bana, Angela Matta, Jeni Melo, Fabienne Läderach, Titus K. Maina, Juliana A. Otieno, Chepsiror Dan, Kibet Kibor, Festus Njuguna, Terry Vik, Ann W. Kinyua, Christian Münz, Jeffrey A. Bailey, Ann M. Moormann

**Affiliations:** Department of Medicine, Division of Infectious Diseases and Immunology, University of Massachusetts Chan Medical School, Worcester, MA, USA; Department of Pathology and Laboratory Medicine, Brown University, Providence, RI, USA; Department of Viral Immunobiology, Institute of Experimental Immunology, University of Zürich, Switzerland; Jaramogi Oginga Odinga Teaching and Referral Hospital, Kisumu, Kenya; Moi Teaching and Referral Hospital, Eldoret, Kenya; Department of Pediatric Hematology/Oncology, Indiana University School of Medicine, Indianapolis, Indiana, USA; Center for Global Health Research, Kenya Medical Research Institute, Kisumu, Kenya; Department of Molecular Genetics and Microbiology, Duke University School of Medicine, Durham, NC, USA

**Keywords:** Burkitt lymphoma, tumor infiltrating lymphocytes, tumor microenvironment, single cell RNA sequencing, Kenya, Epstein-Barr virus

## Abstract

Survival outcomes for pediatric Burkitt lymphoma (BL) substantially vary depending on geography (50-90%), which also serves as a proxy for the prevalence of Epstein-Barr virus (EBV) within the tumors. Although BL is considered an immunologically “cold” tumor with few tumor-infiltrating lymphocytes (TILs), their functional status has not been fully evaluated, especially for EBV-positive disease. Here, we characterize the exhaustion and activation profiles of T cells in the tumor microenvironment (TME) of EBV-positive BL using orthogonal methods, single-cell gene expression analysis, spectral flow cytometry, and immuno-histochemistry staining (IHC). We found that CD8+ TILs displayed a mosaic of immune inhibitory gene expression encoding, *PD1*, *TIGIT*, *LAG3* and *HAVCR2/*TIM3. IHC validated the expression of PD1 and TIGIT on CD8+ TILs, as well as their respective ligands, PDL-1, PVR, and Nectin-2 on malignant B cells. Despite exhaustion-associated signatures, CD8+ TILs retain cytotoxic potential, expressing granules (i.e. Granzyme A, Perforin) and cytokines (i.e. IFNγ) and demonstrate an increased uptake of metabolites such as glucose, arginine, and methionine. In peripheral blood, pediatric BL patients exhibited a significantly higher abundance of PD1+TIGIT+ CD8+ T cells compared to healthy children. Notably, these circulating T cells from BL patients express significantly lower levels of TOX, suggesting they are not irreversibly dysfunctional. Together, our results indicate that CD8+ T cells both in the TME and in circulation of children with BL are not terminally exhausted but remain poised for functional re-invigoration. These findings support the potential integration of immune checkpoint inhibitors into combination chemotherapeutic regimens to improve outcomes for these children.

**Significance:** EBV-positive BL tumors contain functional, metabolically active CD8+ T cells.

Circulating PD1+TIGIT+CD8+ T cells found in BL patients’ blood are a biomarker for those in the tumor microenvironment.

## Introduction

Endemic Burkitt lymphoma (BL) is an Epstein-Barr virus (EBV)-associated pediatric cancer,^1^ which occurs more frequently in *Plasmodium falciparum* (Pf) malaria holoendemic areas^2–4^. In these settings, children experience their primary EBV infection before 2 years of age^5,6^ with higher EBV loads^5^, suggesting that chronic Pf infections impair EBV immunosurveillance^7^. Studies have shown that EBV-specific CD8+ T cell responses are essential to clear viral infection but were found to be deficient in children co-infected with Pf-malaria^8–10^, supporting the premise that malaria-induced immune conditioning increases the risk of EBV^pos^BL tumorigenesis^11,12^. More recently, we showed that children with EBV^pos^BL had high frequencies of PD1+CD4+ T cells^13^, leading us to explore T cell exhaustion as a mechanism responsible for the loss of control over EBV.

Chronic antigen stimulation has been shown to induce exhausted T cells^14–18^. This led to characterizing the stages of exhaustion from “stem-cell like T exhausted effector cells” (S-Tex) with high potential proliferation (e.g. CD101**^_^**CD69+CXCR3+CXCR5+Blimp1+ID2+); to “progenitor T-exhausted effector cells” (P-Tex) which express immune checkpoint inhibitors (ICI) such as PD1, TIGIT, CTLA4, LAG3 or TIM3, in addition to Eomes, ID3, Bcl2, Bcl6, CXCR3, Ki67 and CXCR5; to “intermediate or transitory T-exhausted effector cells” (I-Tex) expressing ICIs, KLRG1, Ki67, CCR5, CXCR3, Blimp1, ID2 and NFAT; and finally to the “terminally T-exhausted cells” (T-Tex) which are now referred to as late-exhausted CD39+CD101+ cells also expressing high ICIs, CD69, CD73, CCR5^18–22^. Combined, these studies define T cell exhaustion as a complex phenomena composed of a constellation of inhibitory receptor expression accompanied by a progressive loss of effector functions, loss of proliferation potential, unresponsiveness to homeostatic cytokines, and low metabolism (suppression of glycolysis)^18,23^. Thus, characterizing T cells as completely ‘exhausted’ has become more complex, with some markers originally used to identify a state of “exhaustion” later found to be expressed by functional T cells after activation depending on their context. In fact, the up-regulation of the transcription factor TOX with a decrease of TCF was described as critical for the CD8+ T cell exhaustion program in mice^24^, whereas TOX expression was not limited to exhausted T cells in humans, but found in polyfunctional T cells^25^. Moreover, it was shown that polyfunctional EBV-specific memory CD8+ T cells express TOX with or without TCF^25^. The relative abundance of exhausted T cell subtypes is also essential, as it was shown that ICIs such as anti-PD1, alone or in combination with anti-TIGIT or anti-CTLA4 were more likely to be effective only if the cells were not at the T-Tex stage^26–28^. It would therefore be crucial to assess the T cell exhaustion profile of patients prior to administering restorative immunotherapies.

In this study, we assess the T cell exhaustion profile in children from Western Kenya diagnosed with EBV^pos^BL to determine whether they would be good candidates for ICI therapies. BL is considered a ‘cold tumor’ compared to other B cell lymphomas, however tumor infiltrating lymphocytes (TILs) were sufficiently abundant to interrogate by single cell RNA sequencing and proteomic methods. Interestingly, we found that despite the expression of at least one inhibitory receptor such as PD1, TIGIT or LAG3, the CD8+ TILs were not categorized as T-Tex. Similar observations were made for peripheral blood PD1+TIGIT+ CD8+ T cells using SpectroFlow, as they expressed cytokines upon stimulation with EBV antigens. Finally, we assessed the inhibitory landscape of the tumor microenvironment (TME) to rank the therapeutic potential of PD1, LAG3, and TIM3 pathways in EBV^pos^BL tumors.

## Methods

*(All reagents used are in supp.table 1)*

### Participants

Children diagnosed with BL were enrolled (n=20) from Jaramogi Oginga Odinga Teaching and Referral Hospital (JOOTRH) in Kisumu and Moi Teaching Referral Hospital (MTRH) in Eldoret, Kenya (illustrated in supp.fig 1A). Tumor biopsies (n=6) and peripheral blood (n=15, BL group) were collected prior to receiving cancer treatments. Healthy children were enrolled from malaria endemic Kisumu County (n=15, malaria-exposed [ME]) and low-to-no malarious Nandi County (n=15, malaria non-exposed [M-NE]). Demographic and clinical characteristics are summarized in the supp.table 2. Children who presented with fever or any malaria symptoms were excluded from the study. We selected formalin-fixed, paraffin-embedded (FFPE) tumor sections from a second set of Kenyan BL patients (N=15) as well as non-BL tumor sections (diagnosed as Hodgkin lymphoma [HL], n=13), for immunohistochemistry staining supp.table 3.

### Ethical approvals

The study was approved by the Scientific and Ethics Review Unit at the Kenya Medical Research Institute; the Institutional Research Ethics Committee at MTRH; and from the Institutional Review Board at the University of Massachusetts Chan Medical School. Written informed consent was obtained from each child’s parent or guardian.

### Tumor biopsies

Core needle biopsies were collected from six tumors for scRNAseq (illustrated in supp.fig 1). Patients’ clinical characteristics are summarized in supp.table 4. Tumor biopsies were transported in tissue storage solution and digested with enzymes using Tumor-dissociation kit and MACS-dissociator for 30 minutes at 37°C. Cell suspensions were filtered using 70μm-strainer. Red blood cell lysis buffer was used when needed. Because predominant tumor B cells can mask less abundant TME populations, four tumors were processed by depleting B cells by positive selection. Then, CD19-depleted fractions were frozen in 90% FBS/10% DMSO and placed in liquid nitrogen until use.

**Figure 1:**
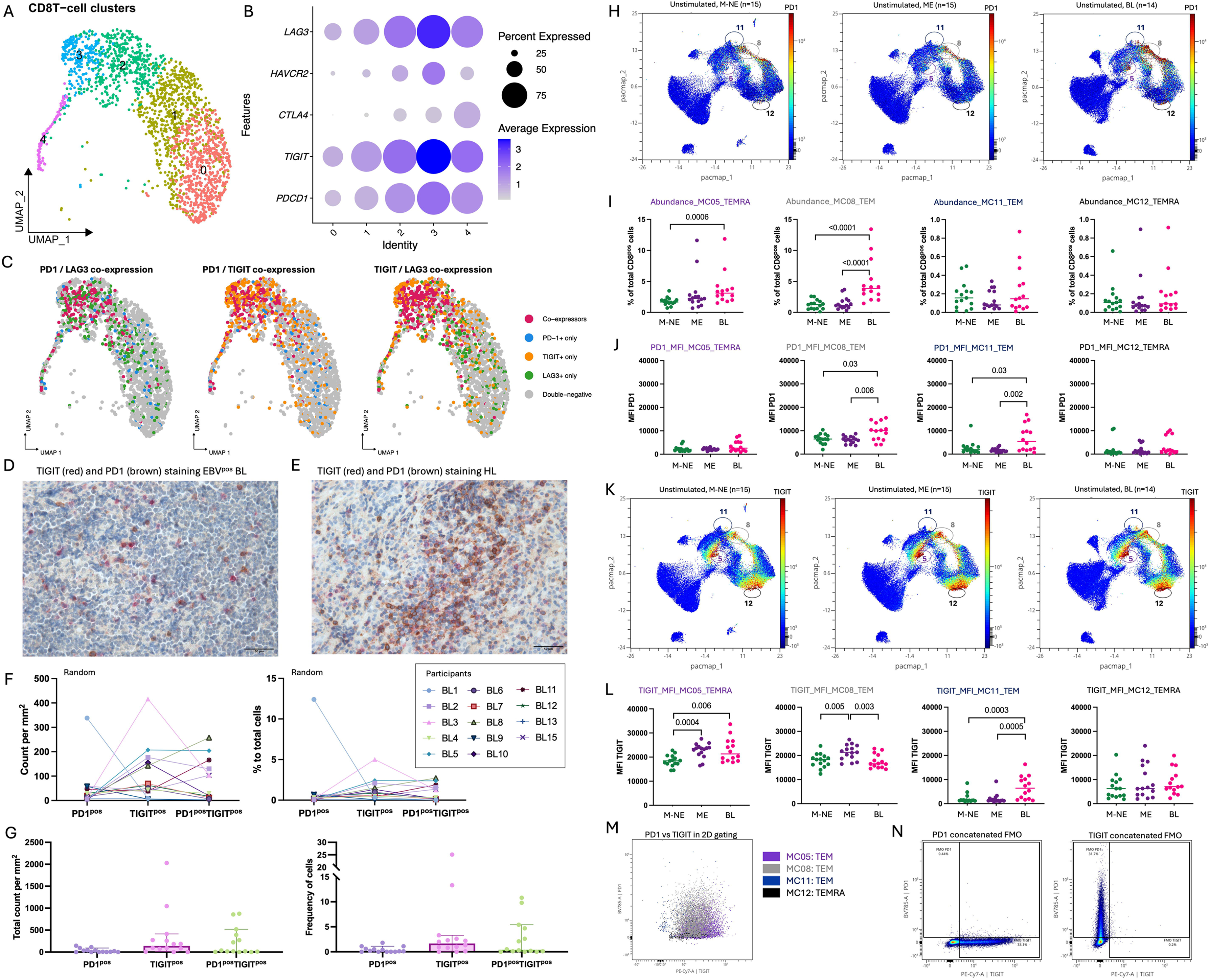
PD1^pos^TGIT^pos^ CD8+ T cells were found in the tumors and peripheral blood of children with BL. **A)** Dimensionality reduction of the RNAseq of the CD8+ TILs clusters (C0 to C4) by UMAP. **B)** Dotplots of the RNAseq of the CD8+ TILs show immune checkpoint inhibitor gene expression within each cluster. Fading blue shades indicate expression level per cell. Dot size indicates frequency of cells expressing the gene of interest. **C)** Overlay of the TIGIT^pos^ (orange), PD1^pos^ (blue), LAG3^pos^ (green), co-expressors (red) and double negative (grey) of the RNAseq of the CD8+ TILs on the UMAP. **D)** Representative IHC image of an BL tumor section where TIGIT is stained in red (E5Y1W antibody developed with new fast red) and PD1 in brown (NAT105 antibody developed with DAB). **E)** Representative IHC image of a Hodgkin lymphoma (HL) tumor section where TIGIT is stained in red (new fast red) and PD1 in brown (DAB). Counts and frequencies of TIGIT^pos^, PD1^pos^, and TIGIT^pos^PD1^pos^ cells across tumor sections of BL patients (n=14) presented by patients **(F)** and altogether (median and 95%CI represented) **(G)**, with the median count of the TIGIT^neg^PD1^neg^ cells being 8,896 cells per mm^2^ (min: 5,822 - max: 12,363) and a median frequency of 97.10% (min: 70.90% - max: 99.80%) of the total cells. **H)** CD8 T cells from peripheral blood samples were compared across our three groups from left to right: malaria non-exposed (M-NE, n=15) and malaria-exposed (ME, n=15) healthy controls and BL cases (n=14). Dimensionality reduction of peripheral CD8^+^ T cells by PacMap where PD1 expression is indicated in color-continued scale. Double positives TIGIT/PD1 CD8 T cell meta-clusters are circled: MC05, MC08, MC11 and MC12. **I)** Abundance of MC05, MC08, MC11 and MC12 within peripheral CD8^+^ T cell subsets of M-NE (green, n=15), ME (purple, n=15) and BL (pink, n=14) children*. **J)** PD1 median intensity expression (MFI) on MC05, MC08, MC11 and MC12 compared between M-NE (green, n=15), ME (purple, n=15) and BL (pink, n=14) children*. **K)** Dimensionality reduction of the peripheral CD8+ T cells by PacMap where TIGIT expression is indicated in color-continued scale. Double positives TIGIT/PD1 CD8 T cells meta-clusters are circled. **L)** TIGIT MFI on MC05, MC08, MC11 and MC12 compared between M-NE (green, n=15), ME (purple, n=15) and BL (pink, n=14) children*. **M)** Overlay of the 4 TIGIT/PD1 double positive meta-clusters on a TIGIT vs. PD1 cytoplot. **N)** Fluorescence minus one (FMO) controls for PD1 and TIGIT staining. *Panels I, J and L were analyzed using a two-tailed Mann-Whitney test where *p*<0.05 is indicated on the plot.

### 10X Single-cell (Sc) RNA-sequencing library preparation

After thawing, viable cells were selected using dead-cell removal kit, counted and resuspended in phosphate-buffered saline (PBS). Sc-suspensions were loaded into the 10x-Chromium Controller, aiming for 10,000 cells, with Chromium-Sc reagent kit following manufacturer’s instructions. Following cell Gel Beads-in-emulsion (GEM) capture and lysis, cDNA was synthesized and amplified to construct sequencing libraries which were sequenced on the Illumina HiSeq 2500 platform at the UMass Chan genomics core. Sequenced reads were processed using the CellRanger toolkit (version 2.1.0).

### Seqwell Sc library preparation

To capture the transcriptome profile of freshly isolated BL tumor cells, and assess whether the TME representation could be obtained without B-cell depletion, we processed two additional BL tumors without B-cell depletion and generated scRNA-seq libraries using SeqWell, a massively parallel, low-input scRNA-seq platform^29^. Briefly, 10,000-15,000 single-cell suspension from each tumor were loaded onto a functionalized polydimethylsiloxane array preloaded with uniquely barcoded mRNA capture beads. The arrays were then sealed with a 10nm pore size hydroxylated polycarbonate membrane, facilitating buffer exchange while confining biological molecules (DNA/RNA) within each well. Subsequent buffer exchange permitted cell lysis, mRNA transcript hybridization to beads, and bead removal before proceeding with reverse transcription. The obtained bead-bound cDNA product then underwent Exonuclease-I treatment to remove excess primer before proceeding with second-strand synthesis, then PCR amplification of the cDNA were purified using the SPRIselect beads, and run on a Fragment analyzer (verification of fragment sizes: 0.7-2kbps). The cDNA libraries were then fragmented and amplified for sequencing using the Illumina Nextera-XT DNA Library kit and custom primers that enabled the specific amplification of only the 3’ ends. The libraries were purified, quantified, and then sequenced on a NextSeq550 system using High output 75 cycle kits at an average of 20,000 reads per cell.

### Single cell transcriptome analysis

Transcriptome data were obtained from 6 BL tumors, a total of 8,007 live cells (mitochondrial gene expression <10%), with a median of 1000 genes detected per cell. scRNAseq libraries from 4 of the BL patients were generated using the 10X Sc platform following negative selection of tumor cells, and the other 2 libraries were generated using Seqwell from freshly collected tumor samples. Demultiplexed FASTQs were aligned to the human reference genome (GRCh38) using STAR aligner^30^. Following alignment, reads were grouped by the 12bp cell barcodes and subsequently collapsed by the 9bp UMI to generate a gene expression matrix. With the gene expression matrix, data normalization (using the variance stabilizing transformation implemented as sctransform in seurat), clustering, and differential expression were performed in R (v.4.0.3) using the Seurat package (v.4.0.4) ^31^. To perform a comparative analysis of the TME while mitigating platform and sample-specific technical variation, we integrated the datasets from the two scRNAseq platforms by merging expression matrices across all cells and applying batch correction and normalization prior to downstream clustering and differential expression. Removal of batch effects arising from differences in library preparation (10X vs SeqWell) improved integration of cells across samples, as demonstrated by visualization on a uniform manifold approximation and projection for dimension reduction (UMAP) plot (supplemental figure 2A), and principal cell-state assignments were consistent across platforms. We attributed clusters to their putative identities and hierarchical similarities using canonical cell markers, corroborated by the top 20 upregulated genes [according to average log fold change (FC) and smallest adjusted P-value] for each cluster compared with the rest of the dataset (supplemental table 5).

### METAFlux analysis of CD8+ T cell subclusters

Metabolic fluxes were inferred using the METAFlux R package following the published workflow^32^. To reduce the impact of sparsity in sc-expression data, we generated cluster-level “pseudo-bulk” expression profiles by bootstrap resampling. For each CD8+ subcluster, cells were sampled with replacement for 100 iterations, and the mean expression of each gene was calculated for each resample. These bootstrap-derived mean-expression profiles were then used to compute reaction activity scores based on gene-protein-reaction rules and to estimate subcluster flux solutions using the observed CD8+ subcluster fractions and a defined nutrient availability constraint (“medium”). Because samples were derived from primary human tumors and matched metabolomics were not available, fluxes were computed using the built-in human_blood medium to approximate physiologic extracellular nutrient availability. METAFlux outputs were reported as relative flux/uptake scores (model-inferred rather than absolute rates). For visualization, flux scores were cube-root transformed, and uptake reactions were sign-adjusted so that higher plotted values indicate higher predicted uptake.

### Immunohistochemistry (IHC)

IHC was performed on a separate series of BL tumors using serial 2μm-thick tissue sections cut from FFPE tissue blocks. Two-color IHC staining of CD4/CD8, PD1/TIGIT, as well as the single staining of PVR, PDL1, PDL2, and Nectin-2 were carried out by Sophistolab AG with the staining procedure as shown in supplemental table 6. For phenotyping and counting analysis, images were acquired using the Vectra-3 slide imaging system. Whole-slide scans were acquired in brightfield mode using a 10X-objective. Regions of interest were subsequently selected and multispectral imaging (MSI) was performed at 20X magnification. Cell segmentation, total cell counts, and phenotype quantification were carried out using the *Tissue Finder Advanced Image Analysis Software*. Individual fields were used to manually train the software for cell segmentation and classification of positive and negative staining, generating classifiers for subsequent algorithmic analysis. The trained algorithm was then applied to the entire section. For the final analysis, only cells classified by the software with a confidence level exceeding 90% were included in the quantification.

### Peripheral blood processing

Peripheral blood was collected in sodium/heparin vacutainers and processed within 2 hours. Briefly, whole blood was centrifuged for 10 mins at 1000g to allow collection of plasma aliquot for serology. Sterile 1XPBS replaced plasma volume for peripheral blood mononuclear cell (PBMC) isolation by Ficoll-paque gradient using Sepmate. PBMCs were suspended in freezing media (FBS, 10%DMSO) and stored overnight at -80°C in Mr. Frosty^TM^ freezing containers. The following day, cryovials were transferred to liquid nitrogen tanks. Samples were shipped to UMass Chan using dry shippers to maintain the cold chain.

### Serology by multiplex beads-based assay (Luminex)

Total IgG antibodies against EBV and *Pf*-malaria were measured using Luminex bead-based suspension assay, as previously described^13^. Previous *Pf*-exposure was determined using recombinant proteins to blood-stage malaria antigens: apical membrane antigen 1 (AMA1), while EBV serology was assessed using the viral capsid antigen (VCA). Briefly, each antigen, or BSA as a background control, was coupled to 5x10^6^ magnetic Luminex microspheres (AMA1 at 5.46 and VCA at 2.5 μg per million beads), and then incubated with study participant plasma (centrifuge at 10,000g for 10mins and diluted 1:100 in assay dilution buffer) for 2hrs, followed by incubation with biotinylated anti-human IgG diluted 1:1000 for 1hr and streptavidin-PE diluted 1:1000 for 1hr following the manufacturer’s instructions. The mean fluorescence intensity (MFI) of each conjugated bead (minimum of 50 beads/antigen) was quantified on a FlexMap3D. Results are reported as antigen-specific MFI after subtracting the BSA value for each individual.

### EBV load by qPCR

DNA was extracted from whole blood cell pellets resuspended in 1XPBS using Zymo kit with the addition of 10mM DTT and Proteinase K during the lysis step. The extracted DNA was then stored at -20°C until use. EBV was detected and quantified by quantitative PCR (qPCR) ^33–35^ using primers and a probe targeting EBV BALF5. Cellular input was quantified in parallel using a human beta-actin reference assay acquired in the HEX channel. Reactions were run in a 12µl total volume containing 6.0µl of 2× iQ Supermix, PCR-grade water (2.5µl), and 2.0µl of template DNA. Final primer/probe concentrations were 500nM each for EBV BALF5 forward and reverse primers (0.6µl each of 10µM stocks), 100nM for the EBV probe (0.12µl of a 10µM stock), and 50nM each for beta-actin forward primer, reverse primer, and probe (0.06µl each of 10µM stocks). qPCR was performed on a Thermal Cycler using the following cycling conditions: 95°C for 3min, followed by 40 cycles of 95°C for 10s and 62.5°C for 30s, with fluorescence acquisition at the end of each 62.5°C step. Quantification was performed using Namalwa genomic DNA standard curves (two integrated EBV copies/cell). EBV standards consisted of six 5-fold serial dilutions spanning approximately 10^5 to 10^1 copies per reaction, run in duplicate. Beta-actin standards consisted of six 3-fold serial dilutions spanning approximately 5×10^5 to 2×10^2 copies per reaction, also run in duplicate. EBV copy number per reaction was interpolated from the EBV standard curve and normalized to cellular equivalents based on beta-actin quantification (assuming two beta-actin copies per diploid cell) to report EBV copies/cell.

### Plasmodium falciparum parasitemia by qPCR

The var gene acidic terminal sequence (varATS) quantitative PCR was used to detect multi-copy genomic sequences as previously described ^36^. Briefly, 0.25μL of PCR water, 6.25μL of 2×LTaqman Universal PCR Mastermix, 0.5μL of 20μM forward and reverse primers, 0.25μL of 20μM probe and 4.75μL of parasite DNA were vortexed and run on a Thermal Cycler with the following cycling conditions: pre-incubation (2min, 50°C), initial denaturation (10min, 95°C), denaturation (15sec, 95°C), and annealing and elongation (1min, 55°C, data collection step). The starting quantity (SQ) values of the parasite samples were estimated against laboratory grown *Pf* 3D7 standard control.

### *Plasmodium falciparum* infected red blood cell lysate (iRBCs)

Synchronized *Pf* trophozoite cultures of iRBCs were isolated using the LD-column and separator stand magnet. After isolation, the purity and number of iRBCs were calculated using a Neubauer cell chamber and Giemsa. Finally iRBCs were frozen in liquid nitrogen for 10mins prior to being placed into a 37°C water bath for another 10mins. This cycle was repeated nine times to ensure good cell lysis. Protein concentration was assessed using the Qubit 4-fluorometer and BR assay. The *Pf*-iRBCs lysate was stored at -80°C.

### *In vitro* stimulation of PBMCs

PBMCs from 15 BL, 15 malaria-exposed and 15 malaria non-exposed children were thawed (using cRPMI and 2mg/ml of DNAse), filtered, and rested overnight at 37°C in a 5%CO_2_ incubator. After resting, cell viability was assessed using Trypan Blue and only samples with equal or greater than 85% viability were used for *in vitro* stimulation (median of 92% viability, min: 85%, max: 98%). One million PBMCs/wells were added to a sterile flat-bottom 96-well plate. EBV peptide pools were added as follows: 2μg/ml of EBNA1 or BLZF1. Stimulation with an equimolar amount of DMSO/PBS as the peptide pool was used as a negative control (<0.1% final), and *Staphylococcus* enterotoxin B (SEB) at 1 μg/ml was used as a positive control. Finally, *Pf*-lysate (5 μg/ml) was added to appropriate wells. Anti-CD28/Anti-CD49d cocktail was added at 1:10 dilution in each well prior to resuspension and incubation for 24 hours at 37°C and 5%CO_2_. GolgiSTOP (7 μg/ml) and GolgiPLUG (5 μg/ml) were added to each well 6 hours prior to the end of the incubation period.

### T cell immunophenotyping by Spectral Flow cytometry

Following *in vitro* stimulation, multiparameter spectral flow cytometry was used to characterize T cells. The following incubations were in the dark. Briefly, cells were washed with 1XPBS and stained with Zombie-NIR for 20mins at room temperature (RT) to exclude dead cells. Cells were then washed with staining buffer (1XPBS, 2% heat inactivated Human serum [HI-HS], 0.02% sodium azide), and incubated for 15mins at RT with 10% HI-HS to block nonspecific binding. Cells were spun for 5mins at 300g and the cell pellet was resuspended in the extracellular antibody cocktail (supplemental table 7) prepared using the stain brilliant buffer, and then incubated for 30mins at 4°C. Cells were then fixed and permeabilized using the transcription factor buffer kit before addition of intracellular antibody cocktail for 30mins at 4°C. Cells were then washed, resuspended in staining buffer and filtered using 70μm-mesh prior being run on Aurora Cytek. Each run included an identical internal control sample to assess variation across the 3 batches and to allow FMOs. Compensation beads were used for live unmixing set-up on the instrument. Median cell viability was 75.5%, ranging from 68.10% to 81.60%.

### OMIQ analysis of spectral flow data

All fcs files were uploaded to the OMIQ platform from Dotmatics (www.omiq.ai). There, scaling and compensation were reviewed and adjusted, if necessary. FlowCut cleaning algorithm was used to remove any aberrant events from the flow files. From all passed events, a median of 218,856 live lymphocytes (min: 66,223; max: 393,694) were recovered. A BL sample was excluded as well as the BLZF1 condition for one malaria non-exposed healthy child, and the iRBCs condition for one BL child due to poor viability and quality report from the FlowCut algorithm. We then subsampled 65,000 live lymphocytes from each fcs file and performed an EmbedSOM dimensionality reduction (supplemental figure 2J-L) to isolate the CD8+ T cell population. Subsampling 5,000 CD8+ T cells per fcs file, we ended up with 220,000 concatenated CD8+ T cells. Using PacMap dimensionality reduction and consensus meta-clustering in FlowSOM (12 meta-clusters), we assessed the expression of each marker within each meta-cluster (supplemental figure 3). Using Prism, we statistically analyzed abundance and marker expression between groups and across stimulation conditions.

### Statistical analysis for flow cytometry

Once the QC, clustering and dimensionality reduction analysis were performed, flow cytometry data was exported in csv. format into GraphPad-Prism (version 10.3.1) for further statistical analysis using Mann–Whitney, Wilcoxon signed-rank, or ANOVA (Friedman for paired data and Kruskal–Wallis for unpaired data), and Dunn’s correction for multiple comparisons. Categorical data, such as sex, was analyzed using a chi-square test. All tests were two-tailed and *p-value*=0.05 was set as the level of significance. Results were expressed either using median with range or the mean with standard deviation (SD).

## Results

### PD1+TIGIT+ co-expressing CD8+ T cells are abundant in the tumor microenvironment and peripheral blood of EBV^pos^BL children

To unravel the function of tumor-infiltrating CD8+ T cells,we assessed the CD8+ T cell exhaustion profiles in BL tumors and peripheral blood from children enrolled in our study. Single-cell RNAseq on six BL tumor biopsies (supplemental figure 2) after CD19 depletion identified the expected malignant B-cell cluster, together with CD8+ T cells, CD4+ Tregs, macrophages, fibroblasts, and plasmablasts (supplemental figure 2A). CD8+ T cells composed 69.9% of the TILs followed by 30.1% of CD4+ Tregs. Natural Killer cells were not detected in the TME. The key gene signatures classifying cell types are indicated in the supplementary figure 2C. Viral gene expression reads for EBER1 and 2 were abundant in the malignant B-cell cluster, further confirming their classification as EBV+ ” (supplemental figure 2B). Focusing on CD8+ T cells, we reclustered them yielding five sub-clusters (clusters 0 to 4, figure 1A). Distinct transcriptional programs are evident across these subclusters, including cytotoxic effector markers (GZMH), activation/exhaustion-associated markers (LAG3/HAVCR2/TNFRSF9), chemokines (XCL1/XCL2/CCL4), and proliferation/cell-cycle genes (MKI67/TOP2A), highlighting functional heterogeneity among tumor-infiltrating CD8+ T cells (supplemental figure 2D). Turning our focus to the expression of selected exhaustion markers *(LAG3, HAVCR2* [Tim3], *CTLA4, TIGIT* and *PDCD1* [PD1]), we found relatively low expression of *CTLA4* and *HAVCR2* (Tim3) within the BL TIL. In contrast, most CD8+ T cell clusters expressed PD1, TIGIT and LAG3 with the highest levels of expression observed in cluster 3 (figure 1B). Because T cell exhaustion is defined by high co-expression of inhibitory receptors, we next assessed exhaustion marker co-expression at the single cell level. Detectable co-expression of multiple exhaustion markers was limited in this dataset, with most CD8+ T cells showing expression of only one marker at a time. Given the sparsity inherent to scRNA-seq data, this likely underestimates the true frequency of co-expression. Nevertheless, CD8+ T cells in cluster 3 showed the clearest evidence of co-expression particularly for TIGIT and PD1 (figure 1C). LAG3 was also co-expressed with other exhaustion markers, predominantly in cluster 3.

To validate the transcription findings, we performed multiplex IHC on an independent set of 15 EBV^pos^BL tumors (supplemental figure 1E). Although overall CD4+ and CD8+ densities were lower among BL tumors than pediatric Hodgkin Lymphoma (HL) (supplemental figure 1F,-I), dual PD1+TIGIT+ cells were detectable in EBV^pos^BL tumors (figure 1D) but to a lesser extent than the HL tumors (figure 1E). Using two analytical methods to quantify mono- or co-expression of PD1 and TIGIT, we found various distributions of exhaustion marker profiles within each tumor (figure 1F, 1G). This analysis revealed that the combined TIGIT+ and double positive TIGIT+PD1+ CD8+ T cells per mm^2^ represented the dominant CD8+ T cell phenotype. Altogether, these results show that CD8+ T cells within the BL TME have heterogeneous exhaustion profiles.

To determine whether this exhaustion phenotype is mirrored in circulating blood, we analyzed PBMCs from EBV^pos^BL patients and compared their CD8 populations to the ones found in healthy children. Using an unsupervised clustering algorithm, FlowSOM (Flow Self-Organizing Map) analysis of CD3+CD8+ events (5,000 cells per participants), identified 12 meta-clusters (MC) (supplemental figure 3). Four MCs that we annotated as TEMRA for terminally differentiated and TEM for effector memory cells (MC05-TEMRA, MC08-TEM, MC11-TEM and MC12-TEMRA) co-expressed PD1 and TIGIT (figure 1H). Two of these (MC05-TEMRA and MC08-TEM) were significantly more abundant in BL patients; MC05 only more abundant in BL compared to M-NE (malaria non-exposed) children (*p*=0.006, figure 1I) while MC08 showed the highest abundance in BL compared to both M-NE and ME (malaria exposed) children (*p*<0.0001 for both, figure 1I). MC08-TEM also showed significantly higher PD1 expression per cells (MFI) in EBV^pos^BL compared to M-NE and ME children (figure 1J, *p*=0.03 and *p*=0.006, respectively). Even though BL children had significantly higher PD1 expression levels, TIGIT levels within this MC were high in ME children compared to both M-NE and BL (figure 1K, *p*=0.005 and *p*=0.003, respectively), with no differences between M-NE and BL children , suggesting an environmental exposure driven TIGIT induction. The other MC of interest was MC11-TEM, which had significantly higher TIGIT expression per cells (MFI) in BL compared to M-NE and ME children (figure 1L, *p*=0.0003 and *p*=0.0005, respectively). The four MCs of interest are shown on the cytoplot of TIGIT vs PD1 (figure 1M), positivity based on the FMOs (figure 1N). These data confirm that BL is characterized by a population of PD1+TIGIT+ CD8+ T cells within the TME that is reflected in their peripheral blood. This supports the collection of peripheral blood as a minimally invasive surrogate to monitor T cell exhaustion within the TME.

### Multiple exhaustion signatures from PD1^pos^TIGIT^pos^ CD8+ T cells

To determine the stage of exhaustion of PD1+TIGIT+ CD8+ T cells in tumors, we assessed markers indicative of activation and differentiation, as T cells transition from pre-exhausted state to terminal exhausted state. From transcriptome analysis, we found cluster 3 also expressed CD44, CXCR3, CD38, HLA-DR (*HLA-DRB1*), high 4-1BB (*TNFRSF9*) and OX40 (*TNFRSF4*) (figure 2A) which confirmed their status as mature and activated, but more importantly, cluster 3 also expressed CD69, CD39 (*ENTPD1*), and CD74, characteristic of T cell exhaustion. CD39 is expressed by both S-Tex and T-Tex cells whereas CD69 expression levels are indicative of terminally exhausted T-Tex cells. In our study population, CD69 is expressed by all CD8+ TIL clusters, yet more so for clusters 0 and 1 suggesting that cluster 3 did not reach the stage of terminal exhaustion. Interestingly high expression of perforin 1, granzyme A and granzyme K were found in all the clusters (figure 2B), whereas granzyme B was more restricted (∼50% or less of the cells). Furthermore, IFNγ expression was found in ∼75% of cells in clusters 1 to 4, whereas TNFα and IL10 expression was low. Thus, the transcriptomic signature showing activation, cytotoxic potential indicates that PD1+TIGIT+ BL TILs are not terminally exhausted.

**Figure 2:**
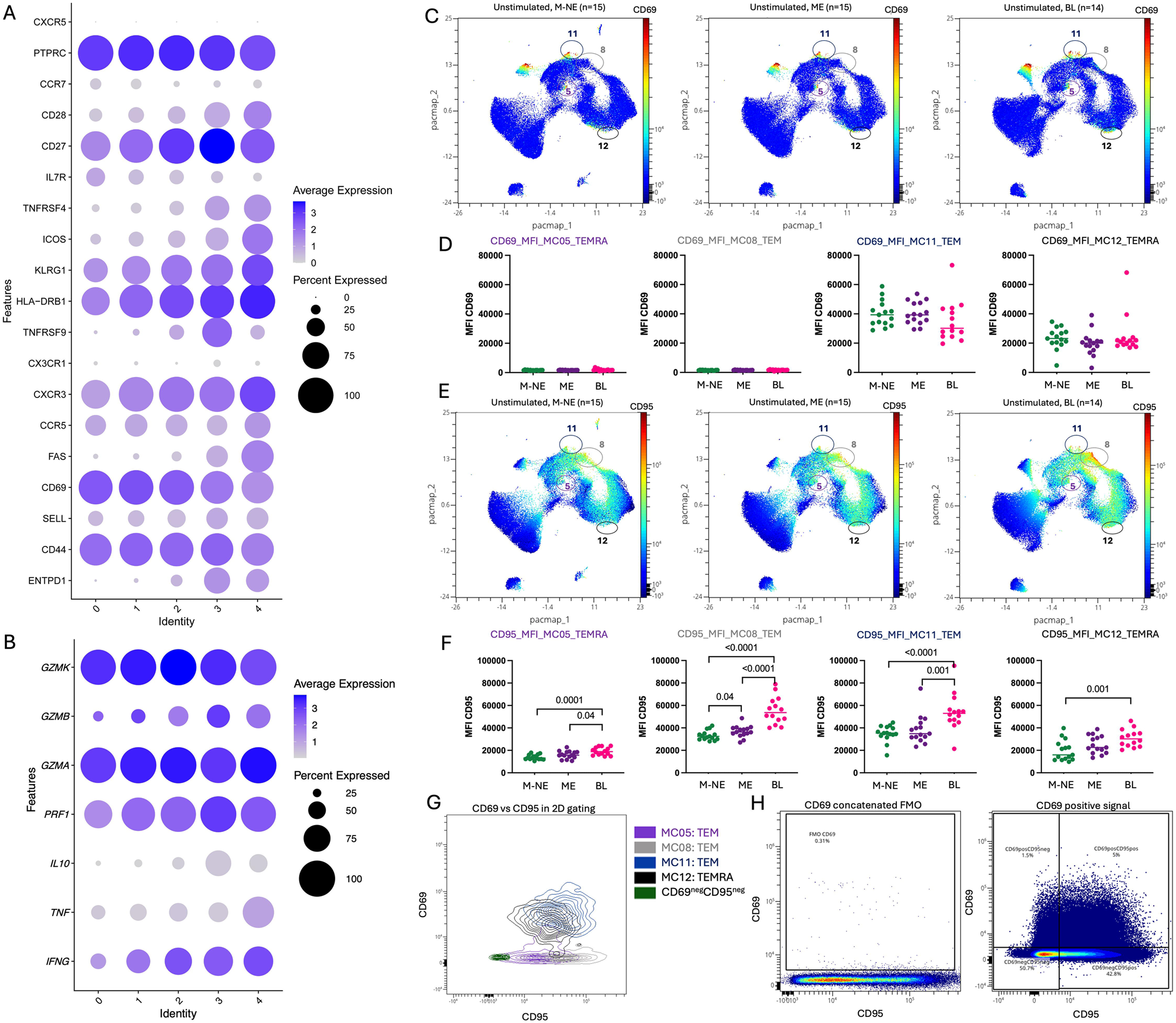
Activation profile of the tumor infiltrating lymphocytes and peripheral CD8 T cells. Dot-plots representing surface markers’ gene expression within each cluster of tumor infiltrated CD8 T cells **(A)** and cytokine (IL-10, TNFα and IFNγ) and granzyme gene expression **(B)** (same metaclusters shown in figure 1A). Blue indicates high expression whereas gray indicates no expression. Bigger the dot is, more cells from the cluster are expressing the marker of interest. **C)** Dimensionality reduction of the peripheral CD8+ T cells by PacMap where CD69 expression is indicated in color-continued scale. Double positives TIGIT/PD1 CD8 T cells meta-clusters are circled. **D)** CD69 MFI on MC05, MC08, MC11 and MC12 compared between M-NE (green, n=15), ME (purple, n=15) and BL (pink, n=14) children*. **E)** Dimensionality reduction of the peripheral CD8+ T cells by PacMap where CD95 expression is indicated in color-continued scale. **F)** CD95 MFI on MC05, MC08, MC11 and MC12 compared between M-NE (green, n=15), ME (purple, n=15) and BL (pink, n=14) children*. **G)** Overlay of the 4 CD69/CD95 double positive cells meta-clusters compared to the CD69^neg^CD95^neg^ population. **H)** FMO control for CD69 staining. *Panels D and F were analyzed using a two-tailed Mann-Whitney test where *p*<0.05 is indicated on the plot.

Finally, proteomic analysis of CD69 and CD95 expression for peripheral PD1+TIGIT+ CD8+ T cells by flow cytometry, found that CD69, involved in T cell migration and retention in tissues was expressed only by MC11-TEM and MC12-TEMRA without differential expression between our study groups (figure 2B-C); whereas CD95 (figure 2D), known to modulate T cell activation relative to the amount of CD95L^37^, was significantly higher for PD1+TIGIT+ CD8+T cells from BL patients compared to M-NE children across the four double positive PD1/TIGIT meta-clusters (MC05-TEMRA *p*=0.0001; MC08-TEM *p*<0.0001; MC11-TEM *p*<0.0001 and MC12- TEMRA *p*=0.001; figure 2E). We observed the same differences compared to ME children for MC05-TEMRA (*p*=0.04), MC08-TEM (*p*<0.0001) and MC11-TEM (*p*=0.001), figure 2F. The four MCs of interest are shown on the cytoplot of CD69 vs CD95 (figure 2G), positivity based on the FMO (figure 2H). Our results suggest that peripheral MC11-TEM in EBV^pos^BL patients might be more prone to be retained in tissues and get activated there (upon low CD95L expression within the TME) compared to PD1+TIGIT+ CD8+ T cells from healthy children.

### PD1+TIGIT+ CD8+ T cells show similar levels of cytokine production with EBV and malaria specific stimulation across groups

To assess EBV antigen specificity of the identified CD8+ T cell subsets, we restimulated PBMCs with overlapping peptide libraries of EBNA1, the only latent EBV protein consistently expressed in BL, and BZLF1, as a representative lytic EBV antigen. From the peripheral blood studies, CD8 effector molecules perforin 1 and granzyme B were only expressed by the MC12- TEMRA subset that also expressed CD69 (figure 3A and 3B, respectively) and unaffected by antigen stimulation. However, 4-1BB expression suggesting T cell activation increased upon BZLF1 stimulation, specifically within the MC11-TEM cells (figure 3C).Using a non-linear dimensionality reduction technique, PacMap, we illustrate the distribution of 4-1BB+ cells across conditions in N-ME, ME and EBV^pos^BL groups (figure 3D-F) showing a significant increase upon BLZF1 for BL children compared to unstimulated cells (figure 3G, *p*=0.006), however, no difference was found across groups (figure 3H). This MC11-TEM effect was confirmed by increased 4-1BB MFI after stimulation (figure 3I) and coincided with higher EBV load in BL patients compared to healthy controls (supplemental figure 4, *p*=0.0003 for both). 4-1BB’s FMO showed in figure 3J. This suggests EBV-specific lytic activation is only seen in BL children expressing PD1 and TIGIT dual positivity but not other groups.

**Figure 3:**
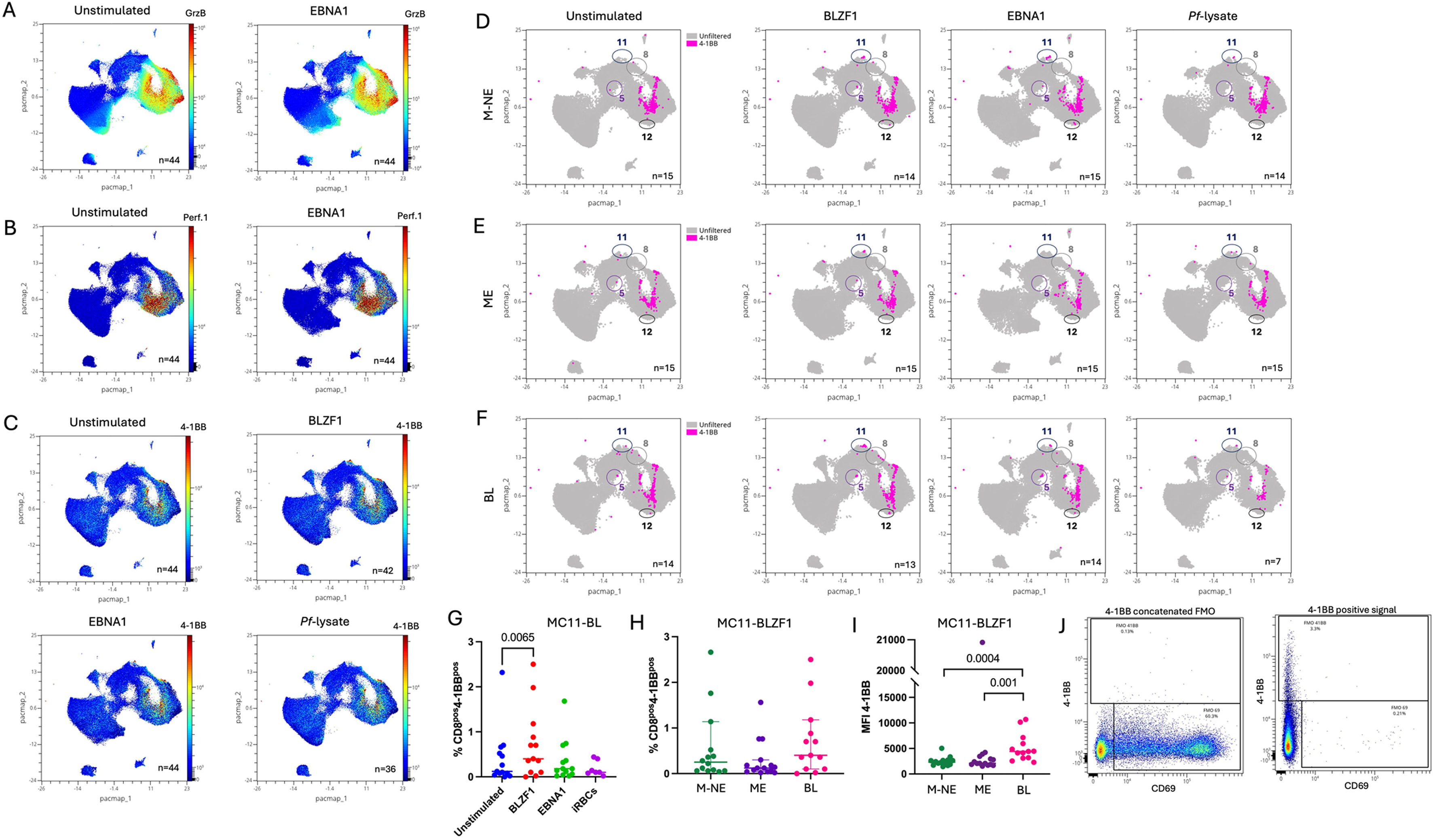
Tumor infiltrating CD8+ T cell clusters express cytokines and granzymes while peripheral TIGIT^pos^PD1^pos^ CD8+ T cells have higher 4-1BB expression upon stimulation in BL patients. Dimensionality reduction of the peripheral CD8+ T cells by PacMap where granzyme B **(A)** or perforin 1 **(B)** expression from unstimulated or EBNA1 stimulated are indicated in color-continued scale. **C)** Dimensionality reduction of the peripheral CD8+ T cells by PacMap where 4-1BB expression is indicated in color-continued scale across the different stimulation conditions (i.e. BZLF1, EBNA1, Pf-iRBC lysate). Overlay of 4-1BB expression (in pink), manually gated, across stimulation on the PacMap representation of the peripheral CD8 T cells from M-NE **(D)**, ME **(E)** and BL **(F)** children. MC05, MC08, MC11 and MC12 are circled. **G)** Frequencies of 4-1BB+ CD8+ T cells from total CD8 T cells from BL patients upon BLZF1 stimulation, manually gated across stimulation. Paired Friedman test where *p*<0.05 is indicated on the plot. **(H)** Frequencies of 4-1BB+ CD8+ T cells from total CD8 T cells from our different groups of children: MNE (Green), ME (purple) and eBL (pink). **I)** 4-1BB MFI on MC11 upon BLZF1 stimulation compared between M-NE (green), ME (purple) and BL (pink) children. Two-tailed Mann-Whitney test where *p*<0.05 is indicated on the plot. **J)** FMO control for 4-1BB staining.

To further explore if this MC-11 TEM cluster expresses cytokines, we assessed IFNγ and TNFα cytokine production upon stimulation using manual gating (supplemental figure 5). An overlay of the IFNγ+, TNFα+ and double IFNγ+TNFα+ cells is shown across stimulation conditions for M-NE (figure 4A), ME (figure 4B) and BL (figure 4C) children. For M-NE children, BLZF1, EBNA1 and *Pf*-lysate each increased the number of IFNγ+ CD8+ T cells (figure 4D, *p*=0.02, *p*=0.003 and *p*=0.007, respectively) showing good effector responses with all stimulations. For ME children however, only EBV antigens induced an expansion of IFNγ+ CD8+ T cells (figure 4E, *p*=0.007 upon BLZF1 and *p*=0.006 upon EBNA1 stimulation) and only BLZF1 induced more γ CD8+ T cells for BL patients (figure 4F, *p*=0.005). This shows that EBV lytic reactivation in BL patients could be useful in eliciting cytokine responses in PD1+TIGIT+ clusters. Regarding TNFα, we noticed a high background in unstimulated cells leading to fewer statistical differences which includes a slightly but significantly higher percentage of TNFα-expressing CD8+ T cells upon EBNA1 stimulation within the M-NE group (figure 4D, *p*=0.04). A similar observation was made for the BL patients showing a significantly higher percentage of TNFα+ CD8+ T cells upon BLZF1 and EBNA1 stimulation (figure 4F, *p*=0.004 and *p*=0.02, respectively). Fewer IFNγ+TNFα+ CD8+ T cells were found but they significantly increased after EBNA1 stimulation in BL patients (figure 4F, *p*=0.003).

**Figure 4:**
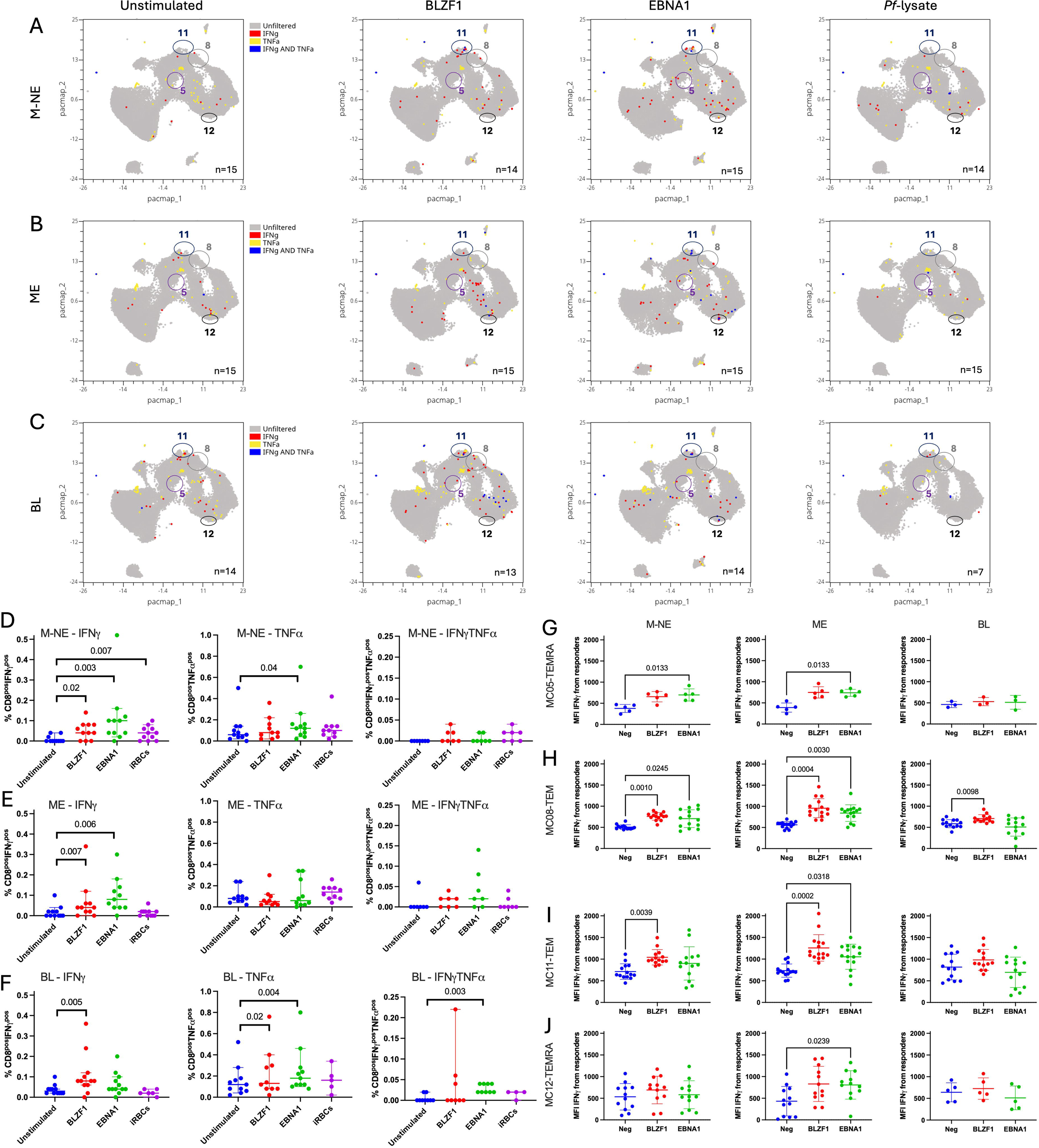
Peripheral TIGIT^pos^PD1^pos^ CD8+ T cells express cytokines upon BZFL-1 stimulation. Overlay of IFNγ^pos^ (red), TNFα^pos^ (yellow), and IFNγ^pos^TNFα^pos^ (blue) expressions, manually gated, across stimulation on the PacMap representation of the total peripheral CD8+ T cells from M-NE **(A)**, ME **(B)** and BL **(C)** children. Double positive PD1/TIGIT metaclusters are circled (MC05, MC08, MC11 and MC12). Frequencies of IFNγ+, TNFα+, and IFNγ+TNFα+ CD8+ T cells, manually gated, from total CD8+ T cells from M-NE **(D)**, ME **(E)** and BL **(F)** children after no stimulation (blue), or stimulation with BLZF1 (red), EBNA1 (green) and iRBCs (purple). IFNγ^pos^ MFI of MC05-TEMRA **(G)**, MC08-TEM **(H)**, MC11-TEM **(I)** and MC12-TEMRA **(J)** meta-clusters across EBV-stimulation from M-NE, ME and eBL children. Panels D to J were analyzed using paired Friedman tests where *p*<0.05 is indicated on the plot.

We confirmed these cytokine responses using meta-cluster analysis. IFNγ-MFI significantly increased upon EBNA1 stimulation within the MC05-TEMRA from the healthy children (figure 4G, *p*=0.01). MC08-TEM showed a significant increase of IFNγ-MFI following BLZF1 stimulation across all participants (figure 4H*, p*=0.001, *p*=0.0004 and *p*=0.009 for M-NE, ME and BL children, respectively), and after EBNA1 stimulation from only the healthy children (figure 4H*, p*=0.02 and *p*=0.003 for M-NE and ME children, respectively). Looking at the overall IFNγ-MFI of MC11-TEM, it was significantly increased upon BZLF1 stimulation (figure 4I*, p*=0.003 and *p*=0.0002 for M-NE and ME children, respectively) but not in BL children suggesting activation without cytokine production (IFNγ); and upon EBNA1 stimulation (figure 4I*, p*=0.03 for ME children) whereas no changes in the MFI were statistically significant across condition within the BL patients’ group. Finally, only EBNA1 stimulation increased the IFNγ-MFI from the MC12-TEMRA from the ME children (figure 4J*, p*=0.02).

In summary, manual gating and meta-cluster analysis corroborate that PD1+TIGIT+ CD8+ T cells can retain cytokine producing capabilities. IFNγ responses were predominant in healthy children, while BL patients displayed notable TNFα responses. Collectively, these results suggest that the PD1+TIGIT+ CD8+ T cells are not terminally exhausted.

### PD1+TIGIT+ CD8+ T cells from healthy children express high levels of TOX upon EBV stimulation

To determine the exhaustion stage within our study population, we examined TOX and TCF, critical transcription factors that regulate CD8+ T cell exhaustion in cancer. TOX, considered a terminal marker of exhaustion in mice^38,39^, has been found in polyfunctional T cells in humans, particularly within EBV-specific CD8+ T cells^25^. Tumor infiltrating CD8+ T cells expressing *BATF* and *Eomes*, but lacking *ILR7* coincide with the initiation of the exhaustion process^15^. Our TIL sequencing data revealed that at least 75% of cells in clusters 3 and 4 express *TOX2*, with more intensive expression in cluster 4 (figure 5A). Transcription factor NR4A, another exhaustion marker interacting with *TOX2* in mouse T cells^24^, was also highly expressed in cluster 3 suggesting that clusters 3 and 4 could be further along in the exhaustion process. However, the lack of terminal exhaustion markers (*CD101)*, low expression of CD39 and the expression of *KLRG1* and *ID2* suggests they are not terminally exhausted but at the intermediate/transitory exhaustion state (I-Tex). In peripheral blood, TOX was not expressed in most of the unstimulated CD8+ T cells (figure 5B to D), but increased after EBNA1 stimulation specifically, and to a greater degree in healthy children. Indeed, EBNA1 induced significantly high TOX expression within all PD1+TIGIT+ meta-clusters across all three groups of children suggesting it’s a response to chronic exposure to EBV (figure 5E to G). The analysis used TOX FMO control (supplemental figure 6A) and this FMO did not change even after EBNA1 stimulation (supplemental figure 6B). An EBV lytic antigen, BLZF1 also induced higher TOX expression, but to a lesser extent (*p*<0.05 for all). Interestingly, TCF expression remained stable regardless of stimulation (figure 5H to J) or group. The PD1+TIGIT+ meta-clusters, notably MC05-TEMRA and MC08-TEM (figure 5H to J), based on the FMOs (supplemental figure 6C), exhibited high TCF expression, suggesting that these cells are not terminally exhausted. Comparing TOX expression across groups of children, TOX was extremely low overall without stimulation (figure 5K). However, BLZF1 induced TOX only within the ME group (figure 5L, *p*=0.0004 and *p*=0.002 compared to M-NE and BL, respectively). This phenomenon became more pronounced upon EBNA1 stimulation where both M-NE and ME children had MC11-TEM cells expressing higher TOX levels compared to BL patients (figure 5M, *p*=0.002 and *p*<0.0001, respectively). Finally, TOX expression did not increase when stimulated with *Pf*-lysate for any of our study participants suggesting there was no correlation with malaria exposure (figure 5N). In summary, even though CD8+ T cells within BL tumors and peripheral blood exhibit some markers associated with exhaustion (i.e. TOX), they are not specific to the BL children group. EBV antigen-specific stimulation showed higher TOX levels within our study and we conclude that their T cells are not terminally exhausted.

**Figure 5:**
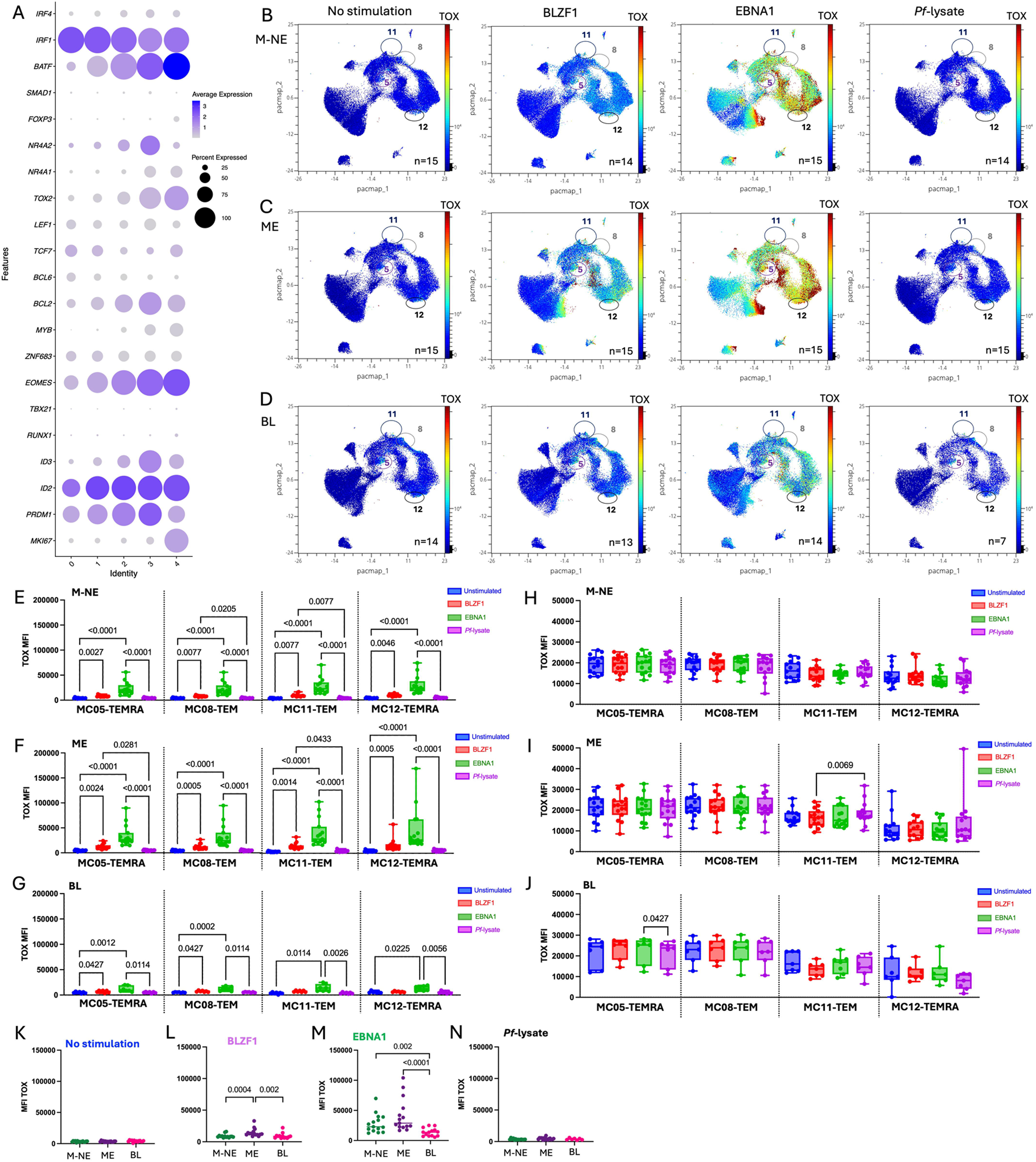
Tumor infiltrating and peripheral PD1+TIGIT+ CD8+ T cells do not express high levels of TOX. **A)** Dot-plots representing transcription factors’ gene expression within each cluster of tumor infiltrated CD8 T cells. Blue indicates high expression whereas gray indicates low expression. Bigger the dot is, more cells from the cluster are expressing the marker of interest. Dimensionality reduction of the peripheral CD8+ T cells by PacMap where TOX expression is indicated in color-continued scale across stimulation conditions (ie. no stimulation, BZFL1, EBNA1 and *Pf*-iRBCs from left to right) within M-NE **(B)**, ME **(C)**, and BL patients **(D)**. Bar-plots of the TOX MFI within each peripheral PD1+TIGIT+ CD8+ T cells meta-clusters from M-NE **(E)**, ME **(F)**, and BL **(G)** upon no stimulation (blue), or stimulation with BLZF1 (red), EBNA1 (green) and iRBCs (purple). Bar-plots of the TCF MFI within each peripheral PD1+TIGIT+ CD8+ T cells meta-clusters from M-NE **(H)**, ME **(I)**, and BL **(J)** upon no stimulation (blue), BLZF1 (red), EBNA1 (green) and iRBCs (purple) stimulations. Dot plot of TOX MFI from the PD1+TIGIT+ CD8+ MC11-TEM meta-cluster across our groups of M-NE (green), ME (purple) and BL (pink) children in absence of stimulation **(K)** or upon BLZF1 **(L)**, EBNA1 **(M)** or iRBCs **(N)** stimulation. Panels E to J were analyzed using paired Friedman tests while panels K to N were analyzed using a two-tailed Mann-Whitney test (*p-*values <0.05 are indicated on the plot).

### Tumor infiltrating PD1^pos^TIGIT^pos^ CD8+ T cells in pediatric EBVpos BL tumors are metabolically active

Terminal exhaustion in T cells is marked by a metabolic shift from glycolysis to dysfunctional oxidative phosphorylation^23^, we assessed the metabolic activity of PD1+TIGIT+ CD8+ TILs using the METAflux pipeline^32^. METAFlux predicted marked heterogeneity in nutrient uptake across CD8+ subclusters (Figure 6). Cluster 3 showed the highest predicted glucose uptake relative to the modeled medium reference (figure 6A), whereas cluster 4 showed comparatively lower glucose uptake, suggesting differences in cell growth and proliferation^40^. In contrast, cluster 4 had higher predicted methionine and arginine uptake (figure 6B and 6C, respectively), nutrients that have been described as key factors in supporting anti-tumor T cell functions^41,42^. Predicted tryptophan uptake scores were low, values under 0, across clusters 2, 3 and 4 (figure 6D), cluster 4 showing the higher range. Tryptophan metabolism is associated with T cell exhaustion as it generates kynurenine^43^, an immunomodulator of T cell functions^44^ which induces PD1 for cytotoxic cells in tuberculosis^45^. Kynurenine uptake scores were low overall but showed a gradual increase from cluster 2 to cluster 4 (figure 6E). Together these findings demonstrate heterogeneity in metabolic potential among CD8+ TILs in pediatric EBV^pos^BL with lower exhaustion potential for cluster 3 showing high glucose uptake and higher exhaustion potential for cluster 4 showing lower uptake.

**Figure 6:**
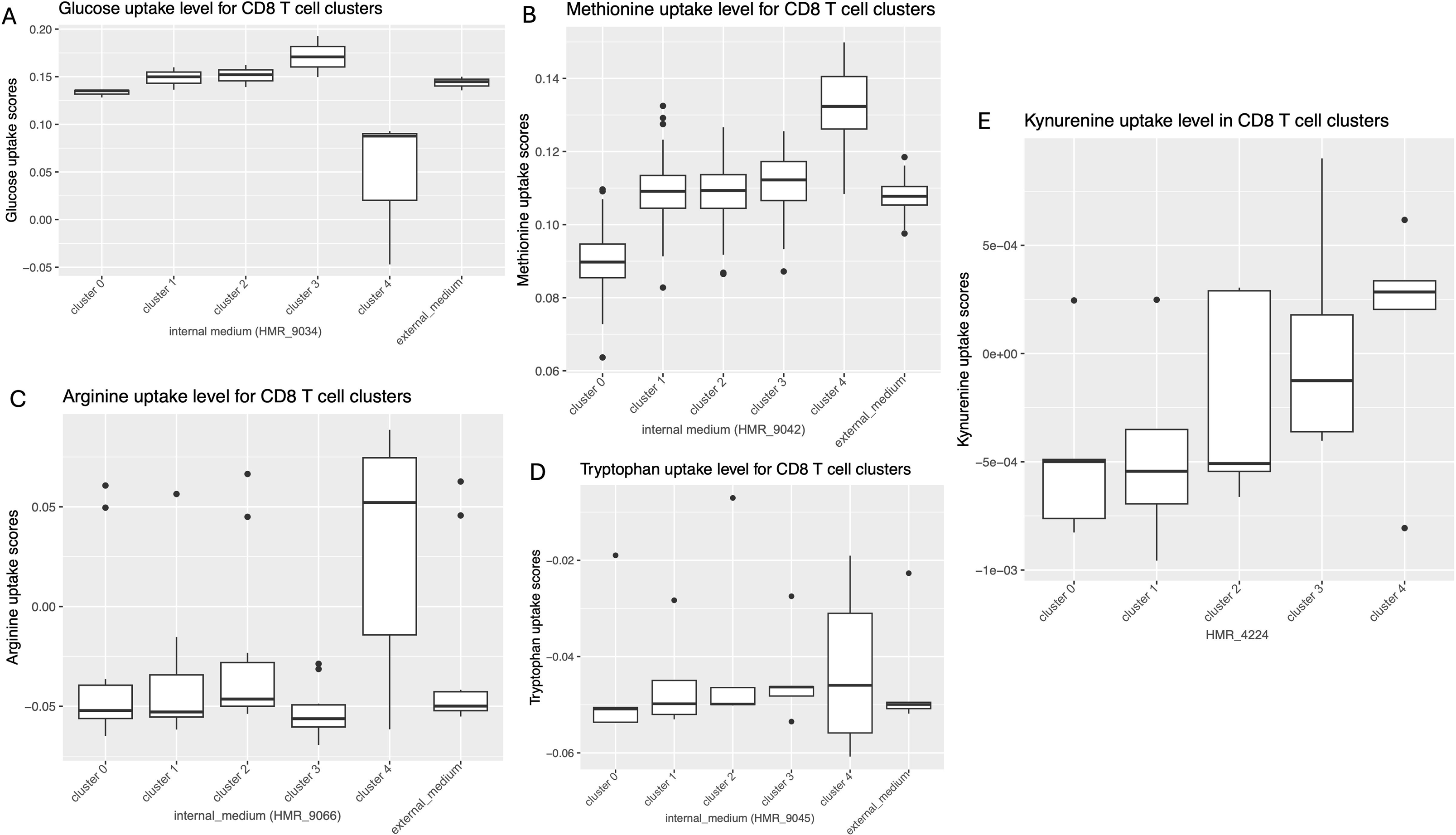
Predicted nutrient uptake programs across clusters of tumor-infiltrating PD1+TIGIT+ CD8+ T cells. Boxplots show METAFlux-inferred relative uptake scores for transport reactions corresponding to **(A)** glucose (HMR_9034), **(B)** methionine (HMR_9042), **(C)** arginine (HMR_9066), **(D)** tryptophan (HMR_9045), and **(E)** kynurenine (HMR_4224) across clusters of tumor infiltrating PD1+TIGIT+ CD8+ T cells. These metabolites were selected because they are linked to T cell activation, function, and responses to the immunosuppressive tumor microenvironment. For each cluster, scores were computed by resampling using 100 bootstraps resamples and visualized as boxplots (the external_medium values are included for reference). Scores represent model-inferred relative flux/uptake under the specified nutrient constraints (human_blood medium), and were cube-root transformed for visualization

### PD1 and PDL1 expression correlate within the BL TME

To further assess the therapeutic potential of ICIs, we next examined cognate ligands expression by tumors using scRNA-seq and IHC. We found that despite the observation that CD8+ TILs expressed high *PDCD1*, few B cell tumors expressed its ligands *CD274* and *PDCD1LG2* genes (figure 7A). However, by IHC, there was a positive correlation between PD1 and PDL1 expression (R^2^=0.59, *p*=0.002), whereas no correlation was found with PDL2 (figure 7B). Pairing TIGIT with its ligands, scRNAseq showed high expression of TIGIT by TILs, but only PVRL4 (nectin-4) was expressed by tumor cells (figure 7C). By IHC, there was no correlation between TIGIT expression with either nectin or PVR (supplemental figure 7). Interestingly, *HAVCR2* (TIM3), was expressed by TILs (figure 7D) but to a lesser extent than the other inhibitory receptors (figure 1B), had high expression of its ligands *PTDSS1* and *HMGB1* by BL tumor cells (figure 7D). Finally, high *LAG3* expression was found within the TILs within only one of its ligands, *CIITA*, being expressed by tumor cells, whereas LGALS3 and SNCA were not detected (figure 7E). These results reveal a varied combination of ICI and cognate ligand pairings for EBV^pos^BL pediatric patients which could guide immunotherapeutic strategies.

**Figure 7:**
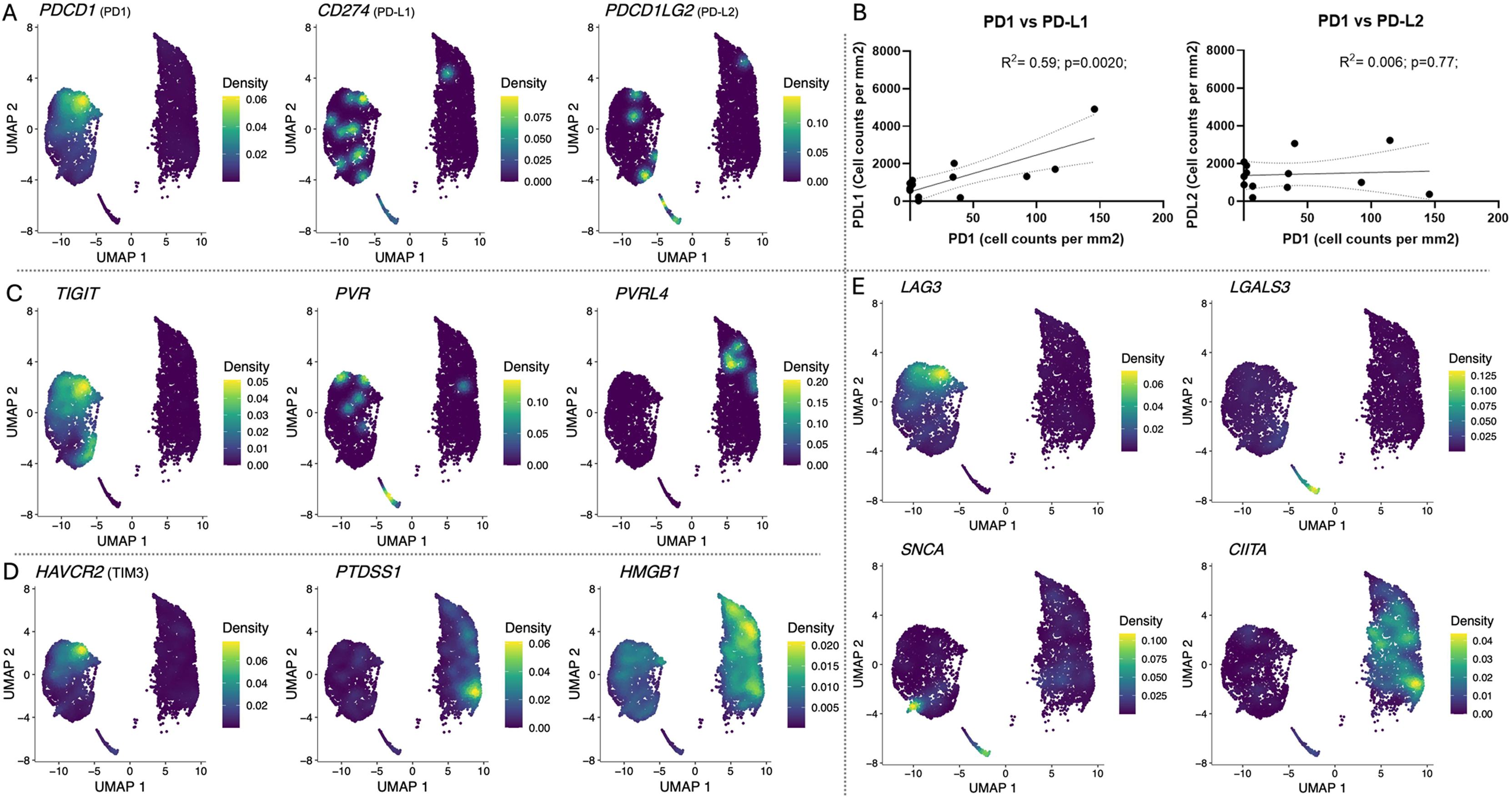
Heterogeneous expression of immune checkpoint inhibitors and their ligands within the BL tumor microenvironment. **A)** PD1 gene expression on the tumor infiltrated immune cells (left cluster) compared to PDL1 and PDL2 gene expression on the tumor cells (right cluster). **B)** Spearman correlation plots of PD1 positive cells vs PDL1 positive cells, and PD1 positive cells vs PDL2 positive cells found in IHC. **C)** TIGIT gene expression on the tumor infiltrated immune cells (left cluster) compared to PVR and PVRL4 gene expression on the tumor cells (right cluster). **D)** TIM3 gene expression on the tumor infiltrated immune cells (left cluster) compared to PTDSS1 and HMGB1 gene expression on the tumor cells (right cluster). **E)** LAG3 gene expression on the tumor infiltrated immune cells (left cluster) compared to LGALS3, SNCA and CIITA gene expression on the tumor cells (right cluster).

## Discussion

T cell exhaustion has been characterized as a loss-of-function that occurs during chronic infections^18^ and cancer^46^. Given the importance of T cells in antitumor immunity, strategies to rescue exhausted T cells have become a major focus of targeted immunotherapies. The goal of our study was to determine the functional state of CD8+ T cells in EBV^pos^ pediatric BL, both in tumors and in circulation. Overall, we found that CD8+ TILs displayed features associated with exhaustion but did not meet criteria for terminal exhaustion (T-Tex) (figure 8), suggesting that these children could still benefit from CD8+ T cell restorative immunotherapies^26,28^.

**Figure 8:**
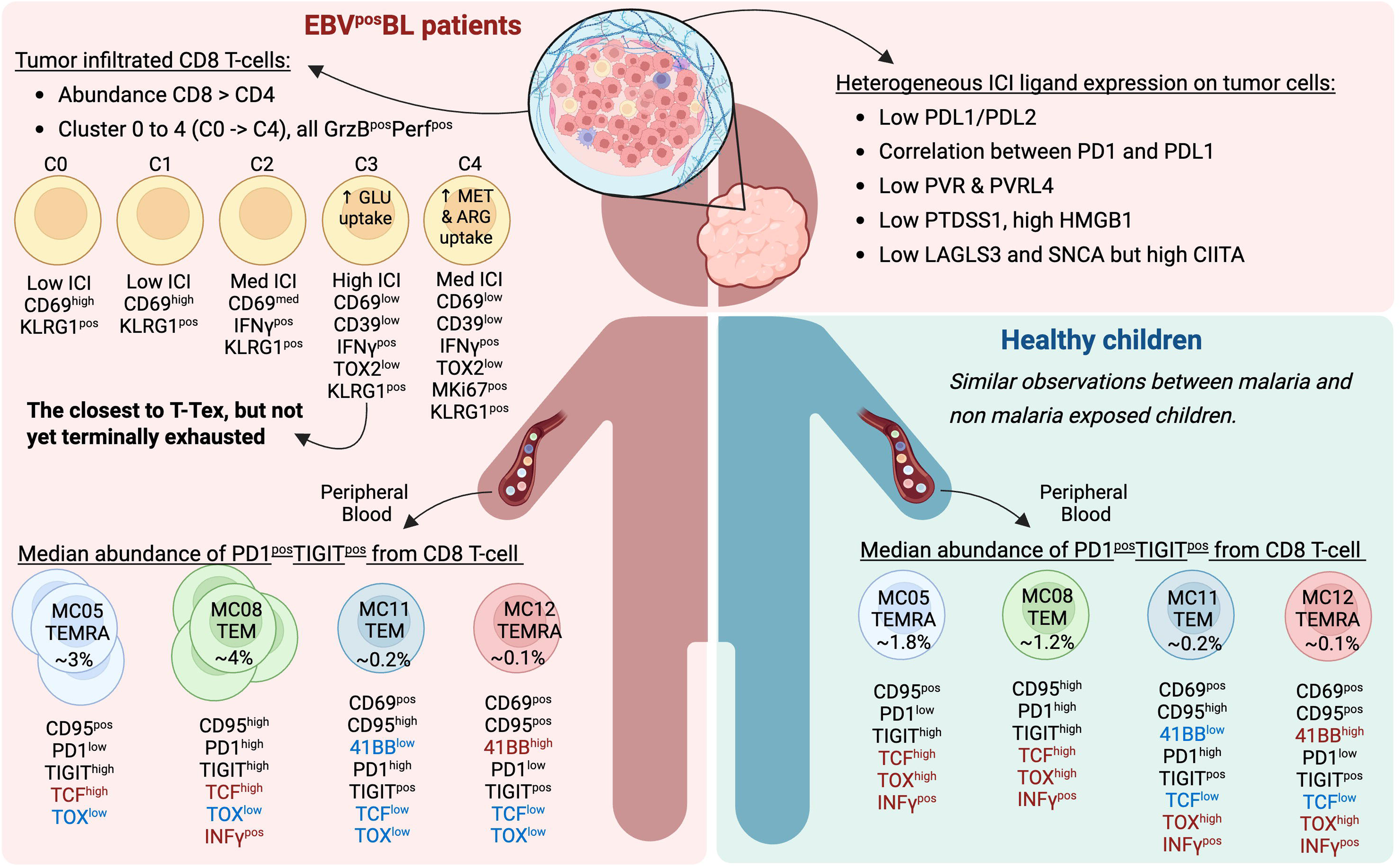
Summary illustration of the differences found in CD8 T cell exhaustion profiles for BL compared to healthy children. Created in BioRender (Agreement number: CA29DHT0EJ)

The most studied ICIs target PD1, TIGIT, LAG3, TIM3 and CTLA4^47,48^ and their accumulation in other cancers is indicative of the degree of T cell exhaustion^18^. In this study of pediatric EBV^pos^BL, we found PD1, TIGIT and LAG3 expression by CD8+ TILs yet only one cluster among five had co-expressing markers reflecting an early stage of exhaustion. Moreover, low frequency of double positive PD1/TIGIT CD8+ T cells were found in peripheral blood of our BL patients and healthy control children (less than 15% of the total CD8+ T cells overall), implying most circulating CD8+ T cells were not on a pathway to exhaustion. There is evidence showing that responses to anti-PD1 therapy in tumors increase the PD1+TIGIT+ circulating population and could be used as an early marker for monitoring therapeutics^49^.

The T cell exhaustion phenotype is not only dependent on ICI expression, but is also characterized by a distinctive transcriptional, metabolic and epigenetic profile that distinguishes them from fully functional, naive or activated T cells^18,23^. Other complementary markers include CD39, CD101, CD69, CD73, and CCR5^18–22^. And yet, our BL tumor CD8+ TILs lacked CD101 and CD73 expression, even though we observed some cells expressing CD39 and CD69. Described as being expressed in the context of HCV and HIV in humans^21^, CD39 is one of several targets against cancer^50^. CD39 is often co-expressed on regulatory T (Tregs) during inflammation, and CD39 and CD73 are enzymes known to transform ADP/ATP to AMP and AMP to adenosine, favoring the establishment of an immunosuppressive tumor microenvironment^51^. CD73 expression was not observed within our PD1+TIGIT+ CD8+ TILs, but it would be interesting to assess both markers on Tregs from EBV^pos^BL patients, as we have previously shown higher frequencies of Tregs in non-survivors^52^. CD69, initially described as an early marker of T cell activation and a tissue retention marker, was more recently described as playing a role in metabolism and regulation of activated tissue resident T cells^53,54^. Interestingly, in our pediatric study population, CD69 expression was found for most PD1+TIGIT+ CD8+ T cells; however, how CD69 regulates anti-tumor immunity and induces exhaustion through CD8+ T cell differentiation is unknown^55^. CD69 expression by CD8+ TILs has been associated with better outcomes in ovarian cancer and melanoma; and with an increase of immune evasion in multiple myeloma as reviewed in^54^. In our study of pediatric BL, we indeed observed CD69 expression in all CD8+ TIL, however the role it plays in patient outcomes remains to be determined.

Reinforcing our first observation that most of the EBV^pos^BL PD1+TIGIT+ CD8+ TILs are not T-Tex, are our metabolic flux results. Indeed, the cluster 3 which had the closest transcriptional signature to T-Tex, showed the highest glucose uptake score. It is well established that CD8+ responses are strongly associated with the glycolytic pathway^56,57^, therefore it is unlikely that CD8+ TIL cluster 3 is T-Tex, despite expressing elevated levels of ICI targeted inhibitory receptors. Other metabolites are also involved in T cell immune responses such as methionine and arginine. Arginine levels can impact proliferation and memory formation of CD8+ T cells preferentially^58^, in T cell epigenetic regulation, and to skew metabolism away from glycolysis^59^. This would explain why cluster 4 (Figure 1A) had low glucose uptake while having high arginine uptake (Figure 6). Cluster 4 also had high methionine uptake, as methionine is involved in remodeling the histone methylation landscape during cell differentiation^42^. Supplementation of methionine in a melanoma mouse model delayed tumor growth^60^, but high levels of SAM (S-adenosylmethionine, produced during the methionine cycle) and MTA (5’methylthioadenosine, part of the methionine salvage pathway), were associated with T cell exhaustion^61^. This duality can be explained by the fact that tumor cells will outcompete T cells for amino acids in the TME while overloading it with MTA and SAM would drive T cell exhaustion^42^. In our study, CD8+ TIL cluster 4 had the highest levels of arginine and methionine uptake suggesting that these cells remain functionally capable. Another metabolic pathway known to influence T cell exhaustion is the tryptophan pathway, leading to the release of kynurenine, an immune modulator of T cell responses^44,45^. We found low tryptophan and kynurenine uptake across our CD8+ TILs, with the exception of cluster 4 utilizing this pathway. To summarize our metabolic flux results, some CD8+ TILs in BL tumors appear to rely on the glucose or arginine pathways to remain functional whereas some may actively contribute to T cell exhaustion.

There are admittedly challenges despite progress made toward introducing ICIs into cancer regimens. Similar to interpatient heterogeneity found in other cancers^62,63^, our results from EBV^pos^BL tumors demonstrate a mosaic of ICI targets within the TME. Therefore, determining the most clinically beneficial combination and dosing strategy would need to be assessed against circumventing inflammatory toxicities^64–66^. Moreover, ICI-based therapies could be combined with anti-CD20 B cell targeting antibodies, which could lead to an increase in EBV-specific memory T cells^67^. Within a clinical trial, if this combination would improve antitumor immunity by targeting EBV^pos^BL tumors warrants investigation.

Our study provides a roadmap for exploring TIL subpopulations in EBV^pos^BL tumors and reveals CD8+ T cells approaching dysfunction but not yet terminally exhausted. Even though BL has been considered a ‘cold tumor’ with fewer TILs compared to other B cell cancers, our study indicates an untapped potential to engage these T cells in anti-tumor immunity. We also found distinct signatures of CD8+ T cell exhaustion in the circulation of EBV^pos^BL patients that also exhibited functionality given their ability to produce cytokines. Studies focused on targeting ICIs trajectory of cells show that PD1 and TIGIT can direct fate of T cells from dLNs to tumor sites with exhaustion progression occurring in tumors ^68^ Determining whether circulating T cells could serve as a liquid biopsy and continue to reflect T cell status within the TME warrant testing within preclinical and clinical trials aiming to improve EBV^pos^BL outcomes.

## Supporting information

Biorender Publication license Fig8

Biorender Publication license supp.fig.1

## Acknowledgements/Funding

The authors would like to thank the children and their families for participating in this study.

This study was funded by NIH, R01 CA189806.

## Authorship contributions

C.S.F: run the flow cytometry- analysis- writing manuscript

C.I.O: run the sequencing- analysis- writing manuscript

P.S.L: design of the assays- analysis- review manuscript

Z.R: run the flow cytometry/qPCR- review manuscript

G.F: run the flow cytometry- review manuscript A.M.: run the serology- review manuscript

J.M.: run the serology/qPCR- review manuscript

1. F. L: run IHC- analysis- review manuscript

T.K.M.: run the sequencing- review manuscript

J.A.O: recruitment patients- review manuscript

F.N: recruitment patients- review manuscript

D.C: collecting tumor biopsies from eBL patients- review manuscript

K.K: run diagnosis of eBL- review manuscript

T.V: recruitment patients- review manuscript

A.K: design of the study- review manuscript

C.M: review all analysis/results- review manuscript

J.B: review the single cells analysis -review manuscript

A.A.M: design of the study- review all analysis/results- review manuscript

## Data availability statement

The single-cell RNAseq data supporting the findings of this study have been deposited in dbGaP under accession number (phs001282.v5). The generated immunohistochemistry and flow cytometry data are available on request.

