## Supplementary material for "Not so cold after all: tumor infiltrating CD8+ T cells in EBV-positive Burkitt lymphoma are quiescent, not exhausted": Supp.Tab.1

|  | **Manufacturer** | **Catalogue Number** | **Item/Instrument/Kit** | **Description/Context** |
| --- | --- | --- | --- | --- |
| **Tumor Biopsies** | Miltenyi | 130-1-00-008 | MACS Tissue storage solution | Transporting tumor biopsies |
|  | Miltenyi | 130-095-929 | Tumor dissociation kit | Enzymatic digestion of biopsies |
|  | Miltenyi | 130-093-235 | gentle MACS dissociator | Mechanical dissociation of tumor samples |
|  | Miltenyi | 130-110-916 | MACS smart strainer (70μm) | Filtering cell suspensions |
|  | Invitrogen | 11143D | Dynabeads™ CD19-PanB | B cell depletion |
|  | Invitrogen | 104-370-28 | FBS (Fetal Bovine Serum) | Freezing media |
|  | Sigma | D2650 | DMSO | Freezing media |
| **10X Single-cell RNA-sequencing library** | Miltenyi | 130-090-101 | Dead Cell Removal Kit | Selecting viable cells post-thaw |
|  | 10xGenomics |  | Chromium Controller | Single-cell RNA-seq |
|  | 10xGenomics |  | Chromium Single Cell 3′ v2 reagent kit | Single-cell RNA-seq |
|  | Illumina |  | HiSeq 2500 platform | Sequencing |
| **Seqwell Single-cell library** | Chemgenes | Macosko-2011-10(V+) | mRNA capture beads | SeqWell platform |
|  | Beckman Coulter | B23317 | SPRIselect beads | cDNA purification in Seqwell prep |
|  | Agilent Technologies |  | Fragment analyzer | Verifying cDNA fragment size |
|  | Illumina | FC-131-1096 | Nextera XT DNA Library Prep Kit | Seqwell library prep |
|  | Illumina |  | NextSeq550 system | High output 75 cycle kits, Sequencing |
| **IHC** | Sophistolab AG (Mutten Switzerland) |  | IHC staining | BL tumor staining |
|  | PerkinElmer |  | Vectra 3 imaging system | IHC image acquisition |
|  | PerkinElmer |  | inForm 2.5.1 Tissue Finder Software | Phenotyping/counting analysis |
| **Blood processing** | BD | 367874 | Sodium/heparin vacutainers | Blood collection |
|  | Invitrogen | 14190250 | 1X-PBS | Blood processing |
|  | Cytiva | 45-001-755 | Ficoll-paque gradient | PBMC isolation |
|  | StemCell Technologies | 85450 | Sepmate tubes | PBMC isolation |
|  | Mr. Frosty |  | Freezing container | PBMC freezing |
| **Serology** | Dr. Sheetji Dutta Walter Reed Army Institute of Research | Gift | Pf-malaria antigen | Atipical Membrane Antigen 1 (AMA1) |
|  | Dr. Japp Middeldorp | Gift | EBV antigen | Vrial Capsid antigen (VCA) |
|  | Sigma | A7888 | BSA | Background control in serology |
|  | BD | 555785 | Biotinylated anti-human IgG | Luminex multiplex assay |
|  | BD | 554061 | Streptavidin-PE | Luminex multiplex assay |
|  | Luminex FlexMap3D multianalyte analyzer |  | Xponent Software | Quantifying bead fluorescence |
| **EBV qPCR** | Zymo Research | D3012 | Quick DNA 96-kit | DNA extraction |
|  | Millipore Sigma | 10197777001 | 10mM DTT | DNA extraction |
|  | Millipore Sigma | P4850 | Proteinase K | DNA extraction |
|  | EBV BLAF5 qPCR forward primer | 5′-CGGAAGCCCTCTGGACTTC-3′ | | |
|  | EBV BLAF5 qPCR reverse primer | 5′-CCCTGTTTATCCGATGGAATG-3′ | | |
|  | EBV BLAF5 qPCR probe | 5′-FAM-TGTACACGCACGAGAAATGCGCCT-BHQ1-3′ | | |
|  | Human beta-actin qPCR forward primer | 5′-TCACCCACACTGTGCCCATCTACGA-3′ | | |
|  | Human beta-actin qPCR reverse primer | 5′-CAGCGGAACCGCTCATTGCCAATGG-3′ | | |
|  | Human beta-actin qPCR probe | 5′-CalFluorOrange560-ATGCCCTCCCCCATGCCATCCTGCGT-BHQ1-3′ | | |
|  | Bio-Rad | 1708860 | iQ Supermix | qPCR reagent |
|  | Bio-Rad |  | CFX96 Real-Time System | C1000 Thermal Cycler, qPCR instrument |
|  | ATCC | CRL-1432 | Namalwa genomic DNA | EBV standards for qPCR |
| **Pf-malaria qPCR** | Applied Biosystems | 4369016 | Taqman Universal PCR Mastermix | Pf varATS qPCR |
|  | varATS qPCR forward primer | 5’-CCCATACACAACCAAyYTGGA-3' | | |
|  | varATS qPCR reverse primer | 5’-TTCGCACATATCTCTATGTCTATCT-3' | | |
|  | varATS qPCR probe | 5’-6-FAM-TRTTCCATAAATGGT-NFQ-MGB-3' | | |
|  | Bio-Rad |  | CFX96 Real-Time System | C1000 Thermal Cycler, qPCR instrument |
| **iRBCs lysate** | Miltenyi | 521-0917-012 | LD column | Isolation of iRBCs |
|  | Miltenyi | 521-0507-256 | Separator stand magnet | Isolation of iRBCs |
|  | Epredia Shandon | 9990700 | Giemsa/Kwik Diff stain | iRBCs counting/staining |
|  | Invitrogen | Q33226 | Qubit 4-fluorometer | Protein concentration measurement |
|  | Invitrogen | A50668 | BR assay | Protein concentration measurement |
| **PBMCs *in vitro* stimulation** | Sigma | DN25 | DNAse | Cell clump treatment |
|  | Invitrogen | 15250061 | Trypan Blue | Cell viability assessment |
|  | Miltenyi | 130-093-613 | PepTivator EBV EBNA-1 | EBV peptide pool for stimulation |
|  | Miltenyi | 130-093-611 | PepTivator EBV BLZF-1 | EBV peptide pool for stimulation |
|  | EMD Chemicals | 324798 | Staphylococcus enterotoxin B (SEB) | Stimulation positive control |
|  | BD | 347689 | Fast-immune anti-CD28 / anti-CD49d | Stimulation |
|  | BD | 554724 | GolgiSTOP | Stimulation |
|  | BD | 555029 | GolgiPLUG | Stimulation |
| **Spectral Flow cytometry** | Biolegend | 423105 | Zombie NIR | Dead cell exclusion - flow cytometry |
|  | GeminiBio | 100-512-100 | Heat inactivated Human serum | Blocking step |
|  | Fisher | S2271 | Sodium azide | Buffer component |
|  | BD | 563794 | Stain brilliant buffer | Flow cytometry |
|  | BD | 562574 | Transcription factor buffer kit | Fixation/permeabilization |
|  | Invitrogen | 01-3333-42 | Compensation beads | Flow cytometry unmixing setup |
|  | Cytek |  | Aurora Flow cytometer (5-lasers) | SpectroFlow® software |
