## Supplementary material for "Not so cold after all: tumor infiltrating CD8+ T cells in EBV-positive Burkitt lymphoma are quiescent, not exhausted": Supp.Tab.2

**Supplemental Table 2: Population characteristics and demographics**

|  | **Malaria non-exposed (M-NE)** | **Malaria exposed (ME)** | **EBV-associated BL (BL)** | **Statistical Analysis** |
| --- | --- | --- | --- | --- |
| **Median Age [range], years** | 8 [5 - 11] | 9 [5 - 10] | 8 [4 - 13] | ^†§^*p*>0.99  ^†^**p*>0.99  ^†#^*p*>0.99 |
| **Sex (% male)** | 6 (40%) | 11 (73%) | 10 (71%) | ^c^*p*=0.11 |
| **Mean hemoglobin (SD), g/dl** | 11.83 (0.90) | 11.52 (1.24) | 9.96 (1.59) | **^†§^*p*=0.004**  **^†^**p*=0.02**  ^†#^*p*>0.99 |
| **Mean WBC (SD), 10^3^/ml** | 5.33 (1.57) | 6.36 (1.76) | 9.63 (5.67) | **^†§^*p*=0.01**  ^†^**p*=0.87  ^†#^*p*>0.99 |
| **EBV qPCR (copies per mg DNA)**  **Median [range]** | 0 [0- 2101] | 0 [0 – 459.8] | 9740 [0 – 1,507,183] | **^†§^*p*=0.0003**  **^†^**p*=0.0003**  ^†#^*p*>0.99 |
| **EBV serology**  **EBNA1 MFI**  **Median [range]** | 82,375  [14,985 – 142,310] | 51,230  [17,617 – 117,495] | 54,425  [20,377 – 129,947] | ^†§^*p*=0.69  ^†^**p*>0.99  ^†#^*p*>0.18 |
| **Malaria qPCR (copies per ml of blood)**  **Median [range]** | 0  [0 – 76.42] | 4.09  [0 – 668.4] | 0.42  [0 – 12,100] | **^†§^*p*=0.04**  ^†^**p*>0.99  **^†#^*p*=0.009** |
| **Malaria serology**  **AMA1 MFI**  **Median [range]** | 2737  [871.5 – 93,640] | 72,635  [7,275 – 136,335] | 41,075  [643.5 – 131,011] | **^†§^*p*=0.02**  ^†^**p*=0.38  **^†#^*p*=0.0001** |

^c^ Statistical test c^2^

^†^ Kruskall-Wallis statistical test with Dunn’s correction for multiple comparison

^§^ Comparison between Malaria non-exposed and EBV-associated BL children

* Comparison between Malaria exposed and EBV-associated BL children

^#^ Comparison between Malaria exposed and Malaria non-exposed children
