## Supplementary material for "Not so cold after all: tumor infiltrating CD8+ T cells in EBV-positive Burkitt lymphoma are quiescent, not exhausted": Supp.Tab.3

**Supplementary Table 3: IHC sample characteristics**

| **Group** | **N** | **Age (median/range)** | **Sex (%Male)** | **Tumor staging (I)** | **Tumor staging (III)** | **Tumor staging (IV)** | **Survival outcome (deceased)** |
| --- | --- | --- | --- | --- | --- | --- | --- |
| EBV^pos^BL | 15 | 9 [3-14] | 80% | 29% | 64% | 7% | 40% |
| HL | 13 | 8 [1-12] | 61% |  |  |  | 33% |
