## Supplementary material for "Not so cold after all: tumor infiltrating CD8+ T cells in EBV-positive Burkitt lymphoma are quiescent, not exhausted": Supp.Tab.4

**Supplementary Table 4: Clinical characteristic of the tumor biopsies.**

| **ID** | **Tumor staging** | **Age (Years)** | **Sex** | **Biopsy (site)** | **Treatment** | **Status** |
| --- | --- | --- | --- | --- | --- | --- |
| EBV^pos^BL-1**^§^** | 3 | 2 | Female | jaw | Received 1 course of treatment. | Deceased |
| EBV^pos^BL-2**^§^** | 1 | 6 | Male | jaw | Finished treatment | Alive |
| EBV^pos^BL-3**^§^** | 4 | 4 | Female | Abdomen | Finished treatment | Alive |
| EBV^pos^BL-4**^§^** | 1 | 12 | Male | Jaw | Relapsed during 5^th^ cycle | Deceased |
| EBV^pos^BL-5***** | unknown | 5 | Male | Jaw | Died before starting treatment | Deceased |
| EBV^pos^BL-6***** | unknown | 9 | Male | Jaw | Died before starting treatment | Deceased |

**^§^** Cryo-preserved tumor single cell suspension sample.

***** Freshly isolated tumor sample
