## Supplementary material for "Not so cold after all: tumor infiltrating CD8+ T cells in EBV-positive Burkitt lymphoma are quiescent, not exhausted": Supp.Tab.5

|  | **p_val** | **avg_log2FC** | **pct.1** | **pct.2** | **p_val_adj** | **cluster** | **gene** |
| --- | --- | --- | --- | --- | --- | --- | --- |
| 1 | 0 | 4.729808702 | 0.977 | 0.875 | 0 | CD8+T | CCL5 |
| 2 | 0 | 3.972606478 | 0.829 | 0.843 | 0 | CD8+T | NKG7 |
| 3 | 0 | 3.864852335 | 0.927 | 0.872 | 0 | CD8+T | GZMK |
| 4 | 0 | 3.706795242 | 0.9 | 0.846 | 0 | CD8+T | GZMA |
| 5 | 0 | 2.920909784 | 0.852 | 0.846 | 0 | CD8+T | CST7 |
| 6 | 0 | 2.465007864 | 0.926 | 0.904 | 0 | CD8+T | CD3D |
| 7 | 0 | 2.25301301 | 0.968 | 0.896 | 0 | CD8+T | IL32 |
| 8 | 0 | 2.17050332 | 0.862 | 0.866 | 0 | CD8+T | HCST |
| 9 | 0 | 1.948877958 | 0.914 | 0.924 | 0 | CD8+T | TRAC |
| 10 | 1.36E-302 | 3.529312812 | 0.76 | 0.77 | 2.72E-299 | CD8+T | CCL4 |
| 11 | 2.46E-173 | 2.188895406 | 0.735 | 0.856 | 4.92E-170 | CD8+T | RARRES3 |
| 12 | 5.22E-162 | 2.115850813 | 0.726 | 0.88 | 1.04E-158 | CD8+T | ITM2A |
| 13 | 6.19E-146 | 2.502899287 | 0.674 | 0.812 | 1.24E-142 | CD8+T | CCL3L3 |
| 14 | 6.07E-141 | 2.702644611 | 0.682 | 0.855 | 1.21E-137 | CD8+T | GZMH |
| 15 | 2.18E-93 | 2.149270957 | 0.631 | 0.8 | 4.36E-90 | CD8+T | PRF1 |
| 16 | 1.65E-58 | 3.326195599 | 0.616 | 0.81 | 3.29E-55 | CD8+T | CCL4L2 |
| 17 | 1.19E-35 | 3.122317301 | 0.348 | 0.762 | 2.39E-32 | CD8+T | GNLY |
| 18 | 6.63E-34 | 2.164026414 | 0.579 | 0.813 | 1.33E-30 | CD8+T | CD8A |
| 19 | 8.29E-14 | 2.440418693 | 0.521 | 0.774 | 1.66E-10 | CD8+T | CCL3 |
| 20 | 3.38E-06 | 2.155770183 | 0.528 | 0.812 | 0.0067565 | CD8+T | IFNG |
| 21 | 1.95E-222 | 1.468590618 | 0.979 | 0.908 | 3.90E-219 | Tregs | IL32 |
| 22 | 8.18E-219 | 1.594101735 | 0.969 | 0.914 | 1.64E-215 | Tregs | TRAC |
| 23 | 6.15E-192 | 1.677275668 | 0.931 | 0.855 | 1.23E-188 | Tregs | CD2 |
| 24 | 4.21E-175 | 1.286351041 | 1 | 0.999 | 8.42E-172 | Tregs | MALAT1 |
| 25 | 2.19E-162 | 1.587386131 | 0.919 | 0.84 | 4.39E-159 | Tregs | JUNB |
| 26 | 3.79E-162 | 1.698487652 | 0.915 | 0.865 | 7.59E-159 | Tregs | S100A4 |
| 27 | 5.28E-144 | 1.341738163 | 0.779 | 0.706 | 1.06E-140 | Tregs | CORO1B |
| 28 | 2.73E-143 | 1.213588664 | 0.853 | 0.793 | 5.45E-140 | Tregs | EMP3 |
| 29 | 1.29E-139 | 1.257435614 | 0.939 | 0.803 | 2.57E-136 | Tregs | HLA-E |
| 30 | 2.72E-138 | 1.227384271 | 0.96 | 0.865 | 5.45E-135 | Tregs | PTPRC |
| 31 | 1.70E-122 | 1.252833239 | 0.845 | 0.801 | 3.41E-119 | Tregs | TRBC1 |
| 32 | 8.92E-121 | 1.216216589 | 0.825 | 0.74 | 1.78E-117 | Tregs | RGS1 |
| 33 | 9.05E-121 | 1.32462547 | 0.887 | 0.807 | 1.81E-117 | Tregs | S100A6 |
| 34 | 2.79E-116 | 1.819401524 | 0.704 | 0.64 | 5.59E-113 | Tregs | KLRB1 |
| 35 | 4.80E-114 | 1.234385112 | 0.944 | 0.903 | 9.61E-111 | Tregs | SH3BGRL3 |
| 36 | 9.05E-110 | 1.549134631 | 0.922 | 0.841 | 1.81E-106 | Tregs | LTB |
| 37 | 2.16E-89 | 1.365441085 | 0.675 | 0.602 | 4.33E-86 | Tregs | IL7R |
| 38 | 6.49E-81 | 1.640746218 | 0.667 | 0.631 | 1.30E-77 | Tregs | TNFRSF4 |
| 39 | 1.51E-31 | 1.235815792 | 0.612 | 0.639 | 3.02E-28 | Tregs | TBC1D4 |
| 40 | 2.47E-13 | 1.464506428 | 0.302 | 0.289 | 4.95E-10 | Tregs | HBB |
| 41 | 2.33E-112 | 6.000507445 | 0.988 | 0.417 | 4.66E-109 | Monocytes/Macrophages | LYZ |
| 42 | 3.85E-110 | 4.227209141 | 0.988 | 0.528 | 7.70E-107 | Monocytes/Macrophages | TYROBP |
| 43 | 1.53E-106 | 3.921875281 | 0.977 | 0.383 | 3.05E-103 | Monocytes/Macrophages | FCER1G |
| 44 | 1.57E-105 | 3.826004793 | 0.977 | 0.463 | 3.13E-102 | Monocytes/Macrophages | CST3 |
| 45 | 1.81E-105 | 4.042512401 | 0.971 | 0.538 | 3.62E-102 | Monocytes/Macrophages | RP11-1143G9.4 |
| 46 | 8.06E-104 | 3.781552672 | 1 | 0.93 | 1.61E-100 | Monocytes/Macrophages | FTH1 |
| 47 | 1.64E-101 | 3.816911085 | 1 | 0.917 | 3.27E-98 | Monocytes/Macrophages | FTL |
| 48 | 1.93E-95 | 3.153563729 | 0.947 | 0.484 | 3.86E-92 | Monocytes/Macrophages | CD68 |
| 49 | 4.04E-95 | 3.845958179 | 0.977 | 0.508 | 8.08E-92 | Monocytes/Macrophages | AIF1 |
| 50 | 2.17E-91 | 3.20913507 | 0.854 | 0.32 | 4.33E-88 | Monocytes/Macrophages | C1QA |
| 51 | 5.96E-91 | 3.454723986 | 0.977 | 0.582 | 1.19E-87 | Monocytes/Macrophages | NPC2 |
| 52 | 4.23E-86 | 3.490310291 | 0.947 | 0.52 | 8.46E-83 | Monocytes/Macrophages | PSAP |
| 53 | 1.19E-85 | 3.048640031 | 0.906 | 0.433 | 2.38E-82 | Monocytes/Macrophages | FCN1 |
| 54 | 3.99E-85 | 2.972443037 | 0.982 | 0.746 | 7.98E-82 | Monocytes/Macrophages | S100A11 |
| 55 | 2.31E-78 | 3.399452751 | 1 | 0.847 | 4.61E-75 | Monocytes/Macrophages | HLA-DRA |
| 56 | 2.40E-78 | 3.092041849 | 0.982 | 0.682 | 4.80E-75 | Monocytes/Macrophages | SAT1 |
| 57 | 7.29E-70 | 4.142998576 | 0.865 | 0.537 | 1.46E-66 | Monocytes/Macrophages | APOE |
| 58 | 2.72E-69 | 4.051795683 | 0.877 | 0.622 | 5.45E-66 | Monocytes/Macrophages | APOC1 |
| 59 | 3.49E-63 | 3.11426996 | 0.883 | 0.587 | 6.99E-60 | Monocytes/Macrophages | CTSB |
| 60 | 1.15E-32 | 3.05037499 | 0.743 | 0.555 | 2.30E-29 | Monocytes/Macrophages | TIMP1 |
| 61 | 0 | 2.824729179 | 0.982 | 0.224 | 0 | Tumor cells | TCL1A |
| 62 | 0 | 2.559186757 | 0.964 | 0.416 | 0 | Tumor cells | CD79B |
| 63 | 0 | 2.432278491 | 0.977 | 0.364 | 0 | Tumor cells | JCHAIN |
| 64 | 0 | 2.139132239 | 0.98 | 0.46 | 0 | Tumor cells | RPL22L1 |
| 65 | 0 | 2.131503067 | 0.807 | 0.216 | 0 | Tumor cells | IGHM |
| 66 | 0 | 2.123084424 | 0.9 | 0.436 | 0 | Tumor cells | HIST1H4C |
| 67 | 0 | 2.073877875 | 0.856 | 0.429 | 0 | Tumor cells | MS4A1 |
| 68 | 0 | 2.059131585 | 0.961 | 0.424 | 0 | Tumor cells | NME1 |
| 69 | 0 | 2.00217396 | 0.934 | 0.397 | 0 | Tumor cells | CD79A |
| 70 | 0 | 1.973455664 | 0.942 | 0.439 | 0 | Tumor cells | HSPD1 |
| 71 | 0 | 1.882552973 | 0.998 | 0.787 | 0 | Tumor cells | YBX1 |
| 72 | 0 | 1.844636407 | 0.982 | 0.584 | 0 | Tumor cells | MIF |
| 73 | 0 | 1.834638096 | 0.93 | 0.434 | 0 | Tumor cells | STMN1 |
| 74 | 0 | 1.82695067 | 0.982 | 0.559 | 0 | Tumor cells | RANBP1 |
| 75 | 0 | 1.801551427 | 0.999 | 0.939 | 0 | Tumor cells | MT-ND4 |
| 76 | 0 | 1.793806912 | 0.99 | 0.567 | 0 | Tumor cells | H2AFZ |
| 77 | 0 | 1.769607871 | 0.952 | 0.506 | 0 | Tumor cells | HMGB2 |
| 78 | 0 | 1.685499711 | 0.999 | 0.845 | 0 | Tumor cells | MT-ND1 |
| 79 | 0 | 1.684558893 | 0.921 | 0.524 | 0 | Tumor cells | HMGA1 |
| 80 | 0 | 1.622139992 | 0.993 | 0.706 | 0 | Tumor cells | EEF1B2 |
| 81 | 5.62E-246 | 1.974669995 | 0.994 | 0.409 | 1.12E-242 | Plasmablasts | LILRA4 |
| 82 | 1.16E-212 | 1.426062095 | 0.991 | 0.355 | 2.31E-209 | Plasmablasts | TSPAN13 |
| 83 | 1.73E-208 | 1.523004592 | 0.98 | 0.234 | 3.47E-205 | Plasmablasts | PLD4 |
| 84 | 3.28E-198 | 2.440707196 | 0.98 | 0.619 | 6.56E-195 | Plasmablasts | IGKC |
| 85 | 4.96E-198 | 1.770338125 | 0.988 | 0.598 | 9.92E-195 | Plasmablasts | RNASE6 |
| 86 | 3.69E-196 | 1.920507889 | 0.997 | 0.576 | 7.37E-193 | Plasmablasts | IRF7 |
| 87 | 3.88E-178 | 1.859924084 | 0.986 | 0.625 | 7.76E-175 | Plasmablasts | GPR183 |
| 88 | 2.92E-146 | 1.397714211 | 0.98 | 0.687 | 5.84E-143 | Plasmablasts | ITM2C |
| 89 | 2.15E-145 | 1.917477597 | 0.997 | 0.67 | 4.30E-142 | Plasmablasts | GZMB |
| 90 | 1.18E-120 | 1.369615972 | 0.96 | 0.574 | 2.36E-117 | Plasmablasts | NPC2 |
| 91 | 6.49E-113 | 1.475229655 | 0.951 | 0.42 | 1.30E-109 | Plasmablasts | TCF4 |
| 92 | 1.32E-105 | 1.412969446 | 0.988 | 0.844 | 2.64E-102 | Plasmablasts | HLA-DRA |
| 93 | 3.37E-102 | 1.69496877 | 0.925 | 0.505 | 6.74E-99 | Plasmablasts | CCDC50 |
| 94 | 1.51E-98 | 1.33215335 | 1 | 0.95 | 3.03E-95 | Plasmablasts | CD74 |
| 95 | 3.03E-92 | 1.714644093 | 0.922 | 0.406 | 6.06E-89 | Plasmablasts | IRF8 |
| 96 | 3.22E-88 | 2.185844269 | 0.859 | 0.472 | 6.44E-85 | Plasmablasts | PLAC8 |
| 97 | 5.11E-72 | 1.66962214 | 0.839 | 0.401 | 1.02E-68 | Plasmablasts | SPIB |
| 98 | 1.73E-71 | 1.65848728 | 0.934 | 0.612 | 3.46E-68 | Plasmablasts | HERPUD1 |
| 99 | 4.79E-70 | 1.575908305 | 0.876 | 0.63 | 9.59E-67 | Plasmablasts | HLA-DMB |
| 100 | 4.44E-45 | 1.390862275 | 0.787 | 0.553 | 8.87E-42 | Plasmablasts | SELL |
| 101 | 2.12E-44 | 5.302482775 | 1 | 0.281 | 4.23E-41 | Fibroblast | CTSK |
| 102 | 1.40E-38 | 4.36371668 | 0.982 | 0.491 | 2.79E-35 | Fibroblast | SPP1 |
| 103 | 1.39E-37 | 2.419728247 | 0.982 | 0.324 | 2.79E-34 | Fibroblast | RAB13 |
| 104 | 2.71E-37 | 4.511848279 | 0.982 | 0.473 | 5.42E-34 | Fibroblast | MMP9 |
| 105 | 3.55E-35 | 4.02274675 | 0.982 | 0.664 | 7.11E-32 | Fibroblast | ACP5 |
| 106 | 8.03E-35 | 2.31748686 | 0.964 | 0.408 | 1.61E-31 | Fibroblast | PRKCDBP |
| 107 | 1.62E-32 | 5.005443542 | 0.964 | 0.471 | 3.25E-29 | Fibroblast | CST3 |
| 108 | 1.71E-30 | 2.481767022 | 1 | 0.81 | 3.43E-27 | Fibroblast | CSTB |
| 109 | 4.37E-30 | 2.463766734 | 1 | 0.588 | 8.74E-27 | Fibroblast | NPC2 |
| 110 | 6.01E-30 | 2.622709195 | 1 | 0.735 | 1.20E-26 | Fibroblast | VIM |
| 111 | 6.68E-29 | 3.399785321 | 0.982 | 0.723 | 1.34E-25 | Fibroblast | ANXA2 |
| 112 | 1.44E-28 | 2.459520462 | 0.982 | 0.611 | 2.88E-25 | Fibroblast | LGALS1 |
| 113 | 5.12E-28 | 2.599191247 | 1 | 0.526 | 1.02E-24 | Fibroblast | PSAP |
| 114 | 9.74E-26 | 2.09517887 | 1 | 0.918 | 1.95E-22 | Fibroblast | FTL |
| 115 | 2.82E-25 | 2.467019117 | 1 | 0.478 | 5.64E-22 | Fibroblast | GPX1 |
| 116 | 4.82E-22 | 2.276953269 | 0.818 | 0.28 | 9.65E-19 | Fibroblast | LUM |
| 117 | 1.84E-21 | 2.342209396 | 0.945 | 0.732 | 3.67E-18 | Fibroblast | CD63 |
| 118 | 3.35E-19 | 2.572920291 | 0.855 | 0.406 | 6.71E-16 | Fibroblast | IFITM3 |
| 119 | 7.88E-06 | 2.439403626 | 0.673 | 0.559 | 0.015764508 | Fibroblast | TIMP1 |
| 120 | 1.30E-05 | 2.405384006 | 0.655 | 0.596 | 0.026075747 | Fibroblast | CALD1 |
