## Supplementary material for "Not so cold after all: tumor infiltrating CD8+ T cells in EBV-positive Burkitt lymphoma are quiescent, not exhausted": Supp.Tab.6

**Supplementary Table 5: Immunohistochemistry list of antibodies**

| **Antibody** | **Clone** | **Company** | **Retrieval (min)** | **Dilution AB** | **Incubation AB (min)** |
| --- | --- | --- | --- | --- | --- |
| CD4 | SP35 | Cell Marque | H2(20) | 1;75 | 30' |
| CD8 | SP16 | Cell Marque | H2(10) | 1;500 | 30' |
| PD1 | NAT105 | Cell Marque | H2(20) | 1;100 | 30' |
| TIGIT | E5Y1W | Cell Signaling | H2(20) | 1;200 | 30' |
| PDL1 | SP263 | Ventana | HIER(48) | Ready to use | 32' |
| PDL2 | 366C.9E5 | Merck | HIER(16) | 1:200 | 26’ |
| PVR | D8A5G | Cell Signaling | H2(10) | 1;100 | 60’ |
| Nectin 2 | D8D3F | Cell Signaling | H2(30) | 1;100 | 60’ |

H2 (Hydrogen Peroxide); HIER (Heat-Induced Epitope Retrieval)
