## Supplementary material for "Not so cold after all: tumor infiltrating CD8+ T cells in EBV-positive Burkitt lymphoma are quiescent, not exhausted": Supp.Tab.7

**Supplemental table 6: Flow cytometry antibodies**

| Marker | Fluorochrome | Company | Cat# | Clone | RRID |
| --- | --- | --- | --- | --- | --- |
| Extracellular cocktail | | | | | |
| CD3 | Spark Blue 550 | BioLegend | 344852 | SK7 | AB_2819985 |
| CD4 | PerCP-Cy5.5 | BioLegend | 300530 | RPA-T4 | AB_893322 |
| CD8 | Pacific Blue | BD Biosciences | 558207 | RPA-T8 | AB_397058 |
| CD95 | PE-Cy5 | BioLegend | 305610 | DX2 | AB_314548 |
| PD1 | BV785 | BioLegend | 329930 | EH12.2H7 | AB_2563443 |
| 4-1BB | BV650 | BioLegend | 309828 | 4B4-1 | AB_2572193 |
| TIGIT | PE-Cy7 | BioLegend | 372714 | A15153G | AB_2632929 |
| CD45RA | BV605 | BioLegend | 304134 | HI100 | AB_2563814 |
| CD69 | FITC | BioLegend | 310904 | FN50 | AB_314839 |
| CCR7 | BV480 | BD Biosciences | 566099 | 3D12 | AB_2739502 |
| Intracellular cocktail | | | | | |
| Granzyme B | APC | BioLegend | 372204 | QA16A02 | AB_2687028 |
| Perforin 1 | Alexa Fluor 700 | BioLegend | 353324 | B-D48 | AB_2750507 |
| IFNg | PE-Dazzle 594 | BioLegend | 502546 | 4S.B3 | AB_2563627 |
| TNFa | BV421 | BioLegend | 502932 | Mab11 | AB_10960738 |
| TCF | Alexa Fluor 647 | BioLegend | 655204 | 7F11A10 | AB_2566620 |
| TOX | PE | Miltenyi Biotec | 130120716 | REA473 | AB_2801780 |
