## Supplementary material for "Not so cold after all: tumor infiltrating CD8+ T cells in EBV-positive Burkitt lymphoma are quiescent, not exhausted": Supp.Captions

**Supplemental tables**

**Supplemental table 1: Reagents**

**Supplemental table 2: Population characteristics and demographics**

**Supplemental table 3: Immuno-histochemistry samples characteristics.**

**Supplemental table 4: Clinical characteristics of the tumor biopsies.**

**Supplemental table 5: Gene signature expression from the tumor infiltrating immune cells.**

**Supplemental table 6: Immunohistochemistry antibodies information**

**Supplemental table 7: Spectral Flow Cytometry antibodies information**

**Supplemental figures**

**Supplemental figure 1: Study design.** Comprehensive schema of the study design, realized with BioRender.

**Supplemental figure 2: Single cell sequencing, immuno-histochemistry and flow cytometry isolation of CD8 T cell population.** **A)** Dimensionality reduction of the EBV^pos^BL tumor cells and tumor infiltrating immune cells using UMAP. **B)** EBER1 and EBER2 gene expression from the single cell sequencing of the biopsies. **C)** Gene signature expression of the different cell types found in the biopsies. **D)** CD8 clusters gene signature. **E)** Representative whole slide scan of tumor section. **F)** IHC staining of EBV^pos^BL tumor section with CD4 (brown) and CD8 (red). **G)** IHC staining of Hodgkin lymphoma tumor section with CD4 (brown) and CD8 (red). **H)** Cell count per mm^2^ of the CD4^pos^ (blue) and CD8^pos^ (red) cells within EBV^pos^BL and HL tumor sections. Mann-Whitney, compare rank, test was applied with FDR of 1% using the two-stage set-up method of Benjamini, Krieger and Yekutieli, *p* value is indicated. **I)** Ratio of CD8/CD4 cells within each type of tumor, EBV^pos^BL (eBL) and HL. Dimensionality reduction of 2.8 million live lymphocytes by EmbedSOM, where CD3 **(J)**, CD8 **(K)** and CD4 **(L)** expressions are shown in colored-continuous scale, red is high and blue is low or no expression.

**Supplemental figure 3: Visualization of the peripheral twelve CD8+ meta-clusters and their protein expression. A)** Dimensionality reduction of 220,000 peripheral CD8+ T cells using PacMap. Each color represents one of the 12 meta-clusters (MC). Colored-continuous expressions of CCR7 **(B)**, CD45RA **(C)**, CD69 **(D)**, CD95 **(E)**, granzyme B **(F)**, perforin 1 **(G)**, TIGIT **(H)**, PD1 **(I)**, TOX **(J)** and TCF **(K)** are shown, red being the highest expression and blue the lowest.

**Supplemental figure 4: EBV viral load.** Dot plot of the EBV copies per μg of DNA from peripheral whole blood from malaria non-exposed (MNE, blue), malaria-exposed (ME, red) and EBV^pos^BL (eBL, green) children. Kruskall Wallis with Dunn’s multiple comparisons test was used, *p* values are indicated.

**Supplemental figure 5: Cytokine cytoplots and manually gated expression across stimulation.** Representative cytoplots of the TNFα versus IFNγ expression from peripheral CD8 T cells across stimulation from MNE **(A)**, ME **(B)** and eBL **(C)**. Frequencies of manually gated TNFα+, IFNγ+, and double positive TNFα+IFNγ+ cells from total peripheral CD8 T cells across stimulation from MNE **(D)**, ME **(E)** and eBL **(F).**

**Supplemental figure 6: TOX and TCF FMOs. A)** TOX FMO in a TOX versus TCF cytoplot upon SEB stimulation. **B)** TOX FMO in a TOX versus TCF cytoplot upon EBNA1 stimulation. **C)** TCF FMO in a TOX versus TCF cytoplot upon SEB stimulation.

**Supplemental figure 7: Correlation analysis between TIGIT and its ligands expression.** Spearman correlation between the count per mm^2^ of positive cells for TIGIT and Nectin-2 (on the left) and TIGIT and PVR (on the right) assessed by IHC.
