## Supplementary figures and images for "Not so cold after all: tumor infiltrating CD8+ T cells in EBV-positive Burkitt lymphoma are quiescent, not exhausted"

### Supp.fig.1

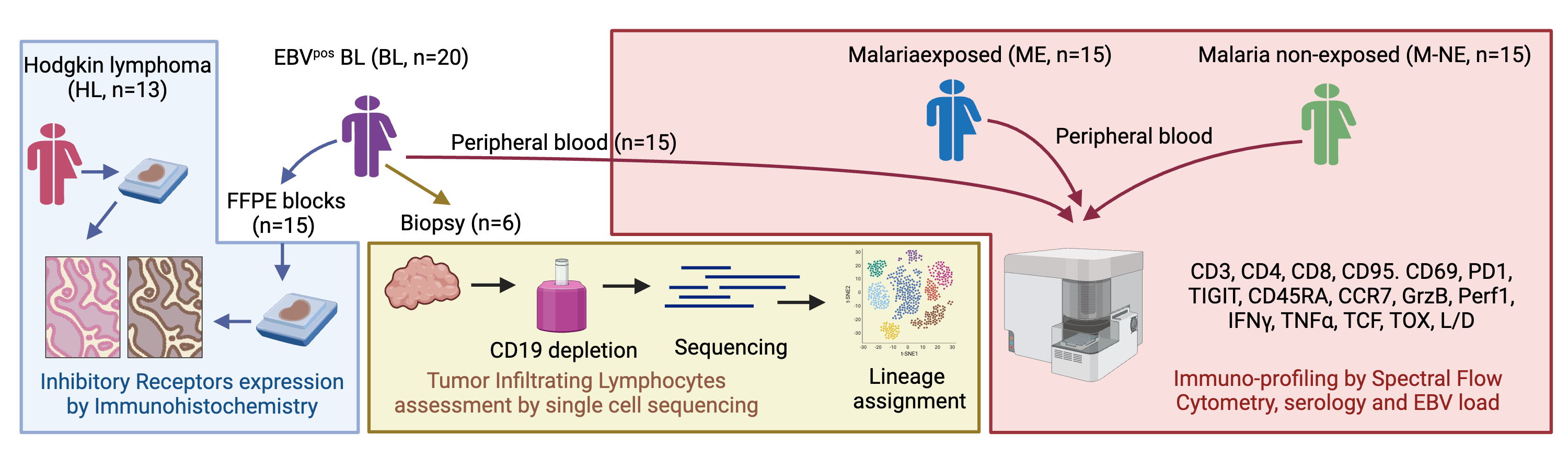

### Supp.fig.2

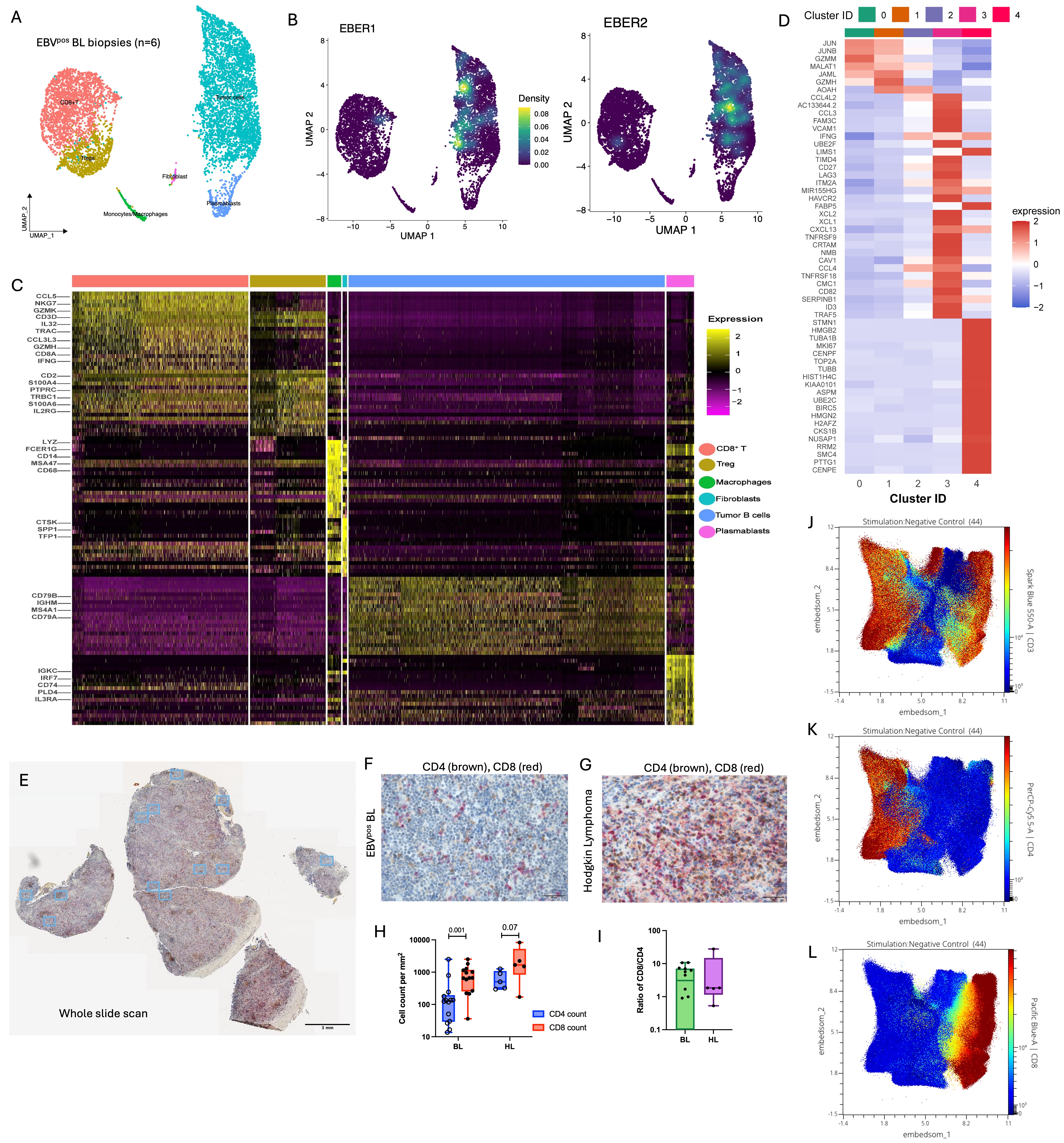

### Supp.fig.3

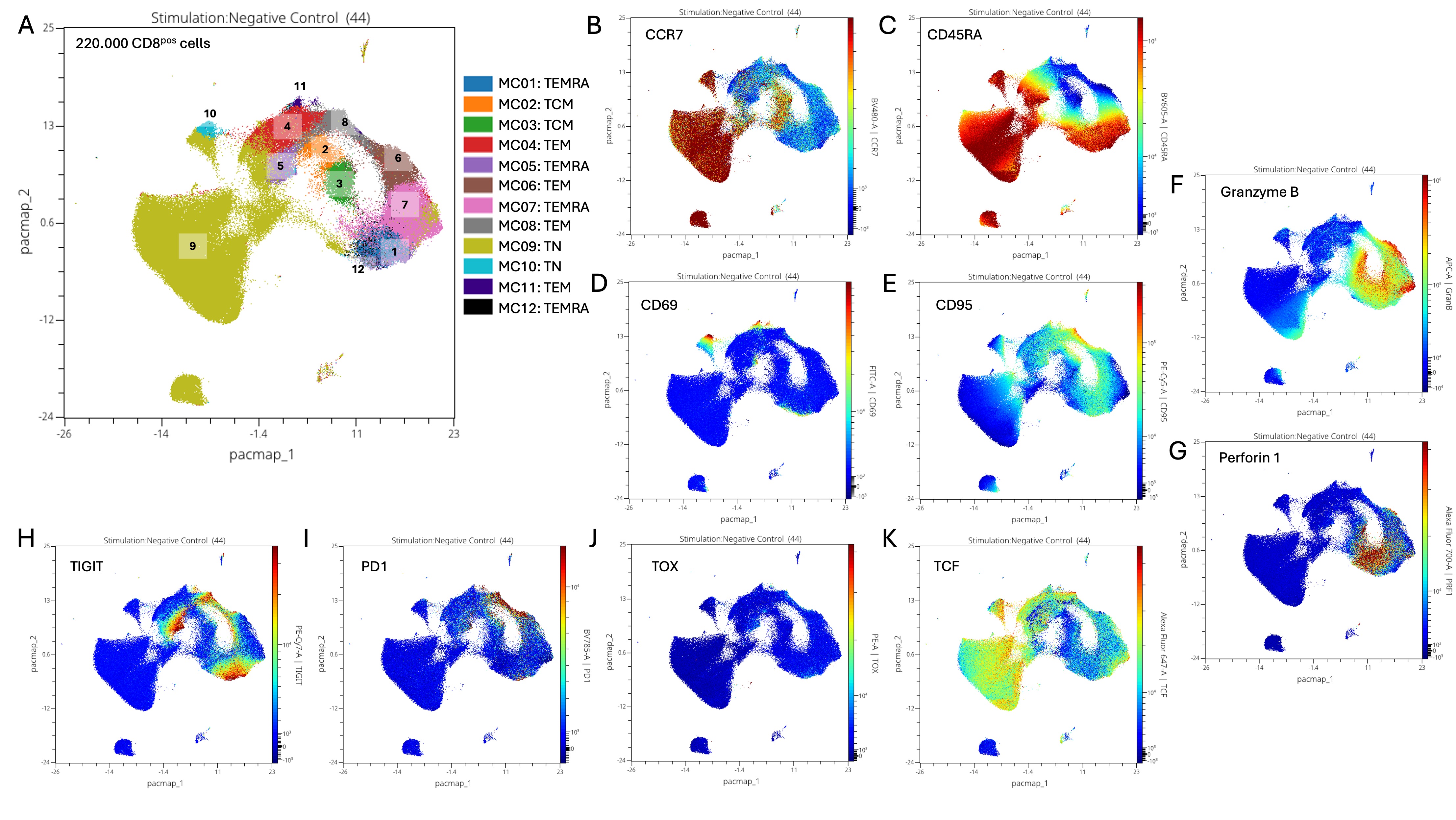

### Supp.fig.4

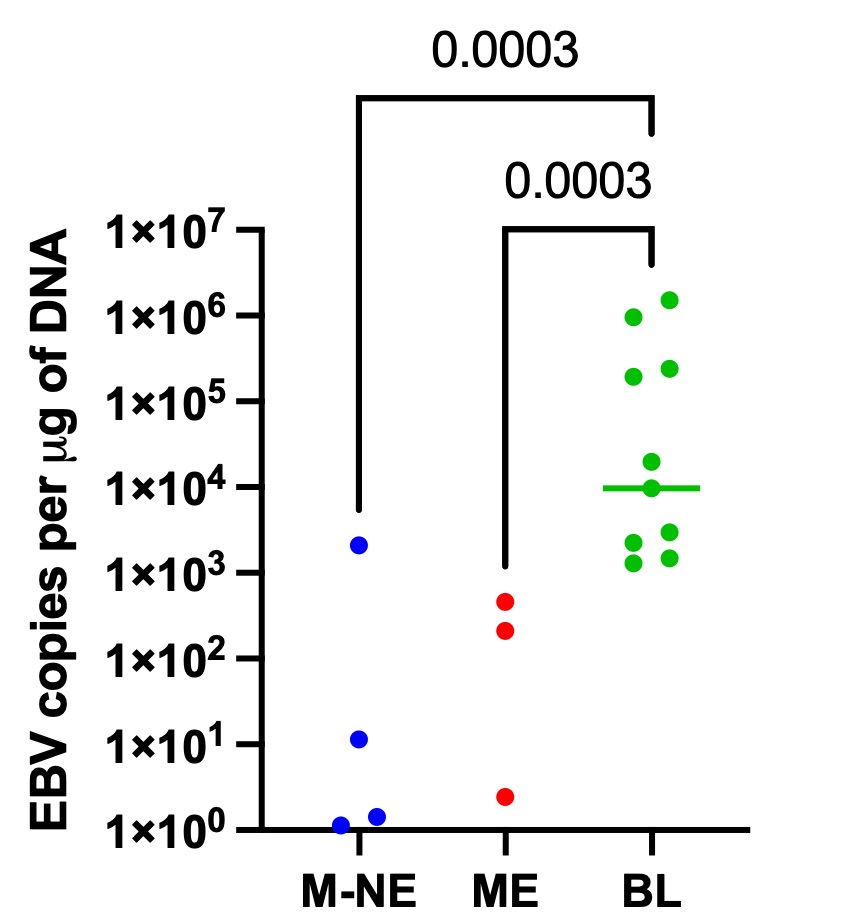

### Supp.fig.5

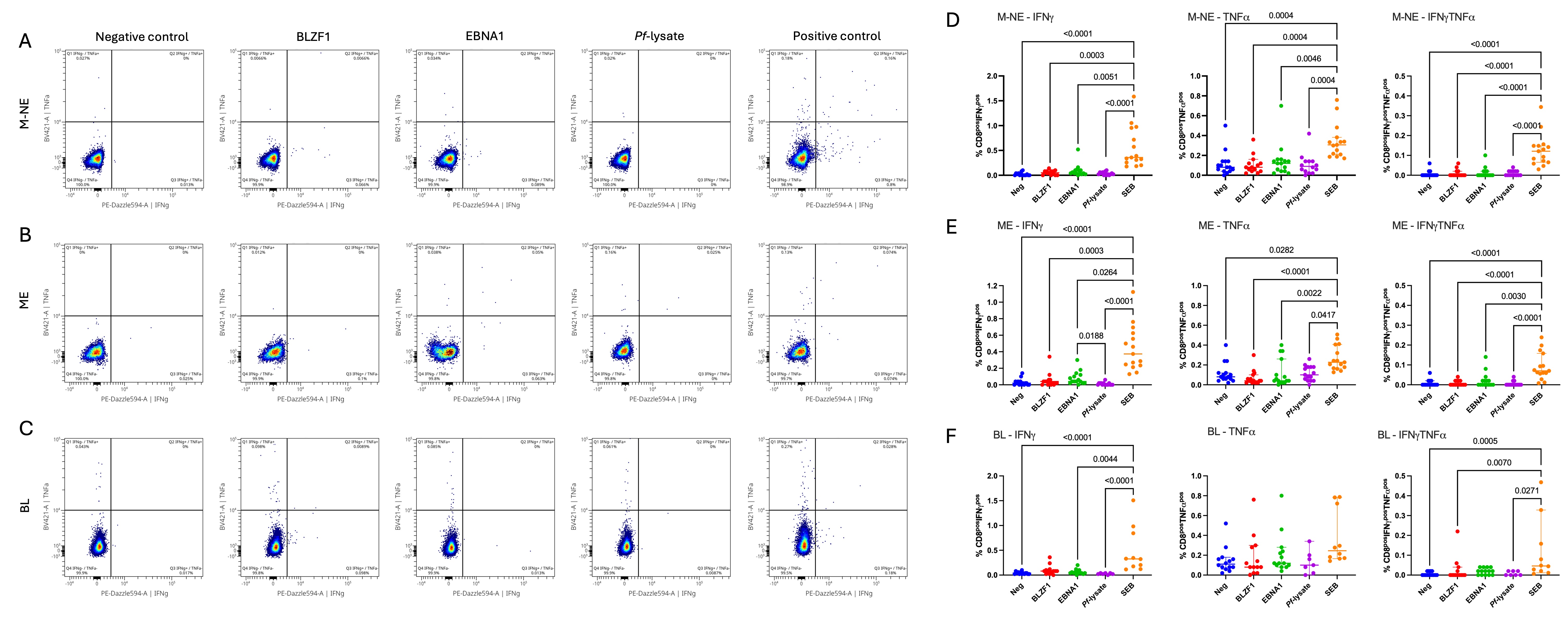

### Supp.fig.6

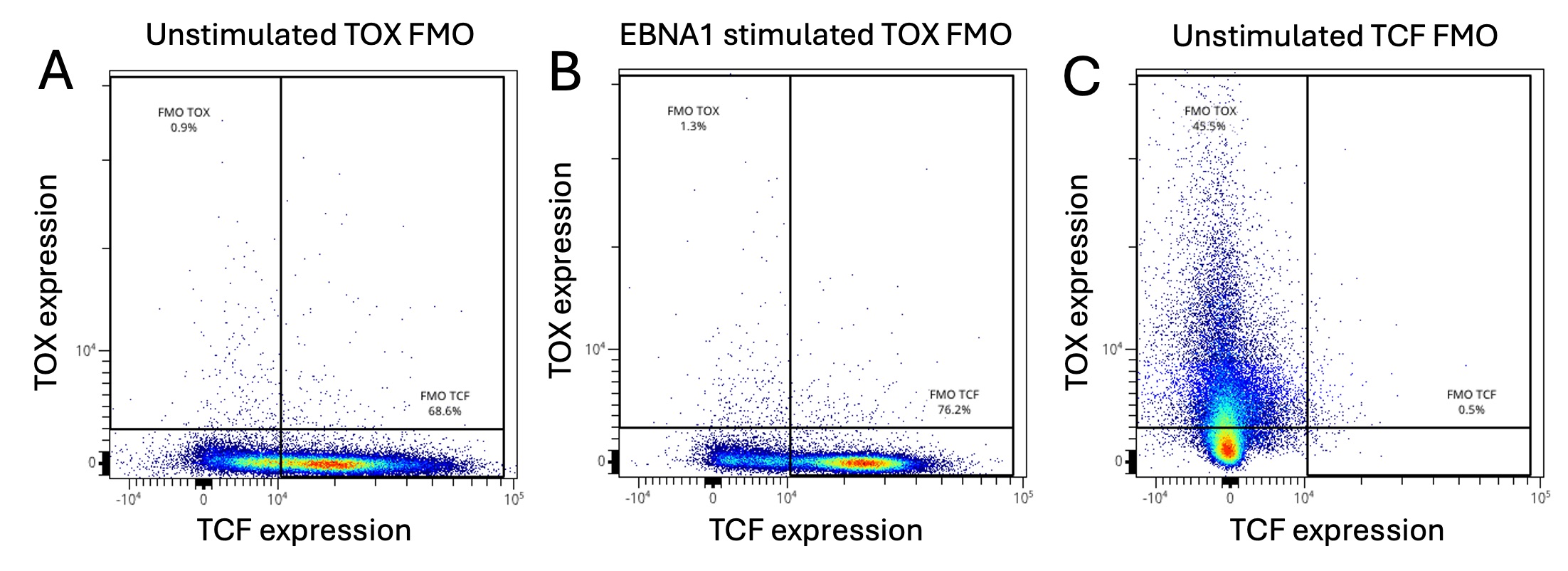

### Supp.fig.7

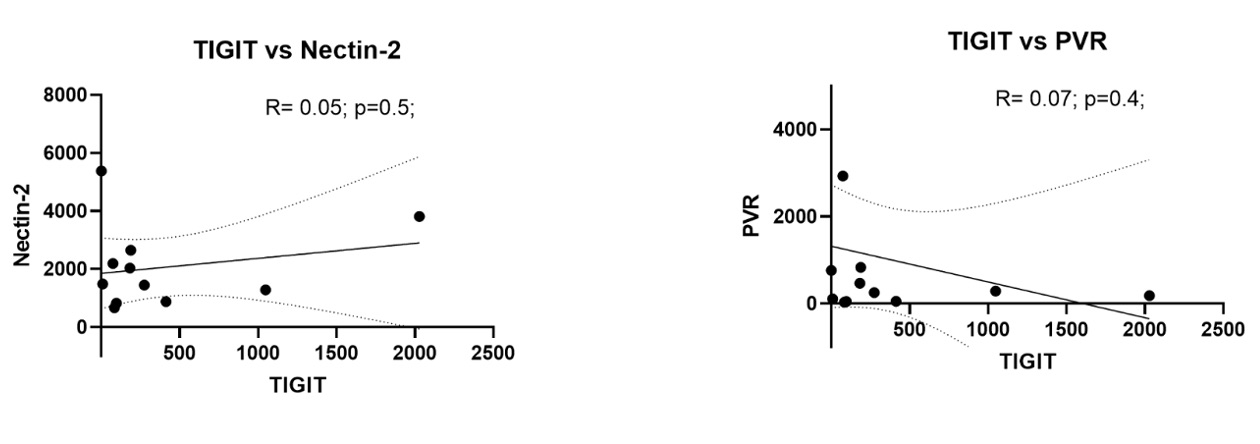
